# ICOS blockade alleviates immune suppression in breast tumor microenvironment

**DOI:** 10.64898/2026.10.22.760908

**Authors:** Shalini Guleria, Chamikara Liyanage, William J Hutchison, Kurtis Treller, Liam Neil, Yingnan Gao, Belinda Yeo, John M Mariadason, Conor J Kearney, Lisa A Mielke, Axel Kallies, Sadanand Vodala, Robin L Anderson, Delphine Merino, Anthony T Papenfuss, Vijay Kumar Yadav, Ajith Vasanthakumar, Stefano Mangiola, Bhupinder Pal

**Affiliations:** School of Cancer Medicine, La Trobe University, Bundoora Vic 3086, Australia; Olivia Newton-John Cancer Research Institute, Heidelberg Vic 3084, Australia; The Walter and Eliza Hall Institute of Medical Research, Parkville, Victoria 3052, Australia; University of Melbourne, Melbourne, Victoria, Australia; South Australian immunoGENomics Cancer Institute, Adelaide, South Australia 5005, Australia; Austin Health, Heidelberg, VIC, 3084, Australia; La Trobe Institute for Molecular Science, La Trobe University, Bundoora, VIC, 3086, Australia; Peter Doherty Institute for Infection and Immunity, Parkville VIC, 3000, Australia; Institute of Molecular Medicine & Experimental Immunology, University Hospital Bonn, Bonn, Germany; Boehringer Ingelheim Pharmaceuticals Inc, Ridgefield, USA; Rutgers New Jersey Medical School, Newark, NJ, USA; Peter MacCallum Cancer Centre, Parkville VIC, 3000, Australia

**Author notes:** These authors contributed equally (Shalini Guleria, Chamikara Liyanage, William J Hutchison). These authors contributed equally (Stefano Mangiola, Bhupinder Pal).

**Keywords:** Breast cancer, metastasis, Tregs, ICOS, immunotherapy

## Abstract

Immune checkpoint blockade has become a standard of care for triple-negative breast cancer, yet only around half of patients benefit. Compensatory immunoregulatory pathways may limit the efficacy of PD-1/PD-L1 blockade. ICOS, an inducible co-stimulatory receptor enriched on activated regulatory T cells (Tregs), represents a potential route of such immune regulation, but its role in breast cancer immunotherapy remains unclear. Here, we identify ICOS as selectively upregulated on tumor-infiltrating Tregs and show that ICOS⁺ Tregs localize within a tumor niche enriched for exhausted CD8⁺ T cells. We confirmed this cellular and spatial association across 1,359 tumor regions from 279 triple-negative breast cancer patients profiled by imaging mass cytometry, and single-cell transcriptomic data from 129 breast cancer donors. Genetic ablation or pharmacological blockade of ICOS disrupted this intratumoral Treg niche, reducing Treg activation and tissue-residency programs while expanding cytotoxic CD8⁺ T cells with a tissue-resident memory phenotype. In syngeneic orthotopic breast cancer models, ICOS blockade restricted primary tumor growth, reduced spontaneous metastasis and prolonged survival without disrupting systemic immune homeostasis. Therapeutic activity was observed with both neoadjuvant and adjuvant treatment, while combined ICOS and PD-1 blockade further enhanced tumor control. These findings support ICOS blockade as a therapeutic strategy to improve checkpoint immunotherapy.

## Introduction

Breast cancer remains the most frequently diagnosed cancer in women worldwide and a leading cause of cancer-related death. Despite advances in early detection and targeted therapies, metastatic and treatment-resistant disease continues to drive poor outcomes. In recent years, immune-based therapies have revolutionized the management of several cancers, with immune checkpoint inhibitors (ICIs) targeting CTLA-4 and PD-1/PD-L1 pathways delivering durable responses and significantly extending survival. However, the success of these approaches in breast cancer has been far more limited [1, 2]. ICIs in breast cancer have shown benefits but mainly confined to a subset of patients with triple-negative breast cancer (TNBC), while hormone receptor-positive and HER2-positive subtypes remain largely unresponsive [3, 4]. This limited efficacy underscores the unique immunobiology of breast cancer, which is generally characterized by a “cold” tumor immune microenvironment with lower levels of T cell infiltration [5, 6].

A major barrier to effective immunotherapy in breast cancer is the profoundly immunosuppressive tumor microenvironment (TME). Within the breast TME, multiple cellular and molecular mechanisms converge to blunt cytotoxic immunity. Tregs, myeloid-derived suppressor cells (MDSCs), and dysfunctional or exhausted cytotoxic CD8^+^ T cells form a suppressive network that prevents effective immune-mediated tumor clearance. Elevated Treg infiltration has been consistently associated with poor prognosis and reduced response to ICIs in breast cancer [7, 8]. Overcoming these suppressive networks is therefore a critical challenge and a prerequisite to unlocking the full potential of immunotherapy across breast cancer subtypes.

Among the immunoregulatory pathways implicated in TME suppression, the inducible co-stimulator (ICOS) receptor has emerged as a key player [9–11]. ICOS, a member of the CD28 family that also includes CD28, CTLA-4, and PD-1, is upregulated upon T cell activation, while its ligand (ICOS-L) is expressed on antigen-presenting cells (APCs) such as dendritic cells, B cells, and macrophages, as well as non-hematopoietic stromal and epithelial cells. ICOS signaling contributes to T cell activation, peripheral tolerance, and immune homeostasis [10] [11]. One of the key roles of ICOS is to sustain the suppressive function of Tregs [12–17]. ICOS⁺ Tregs represent a specialized subset enriched in non-lymphoid tissues such as visceral adipose tissue and mucosal surfaces, where they maintain tissue integrity and limit inflammation [14] [18] [19] [20]. The same regulatory function, however, is exploited by tumors, as ICOS⁺ Tregs accumulate in the TME where they suppress cytotoxic CD8⁺ T cell and natural killer (NK) cell activity, thereby promoting immune evasion [9, 21–23] [24]. Elevated ICOS⁺ Tregs have been reported in melanoma [10], gastric [25], and ovarian [26] cancers, where their expansion correlates with poor prognosis and therapeutic resistance [25, 26]. However, ICOS expression extends beyond Tregs to effector and memory CD4⁺ and CD8⁺ T-cell subsets, suggesting that its function within the breast TME may be context dependent. It therefore remains unclear how ICOS coordinates the spatial organization and suppressive phenotype of tumor-infiltrating Tregs, and how these changes influence effector T-cell responses, primary breast tumor growth, and metastatic progression. Furthermore, previous ICOS-targeting studies have not evaluated orthotopic mammary tumor models capable of spontaneous metastatic dissemination, nor survival outcomes following ICOS blockade. We therefore assessed how ICOS blockade impacts Treg-mediated suppression and effector CD8⁺ T cell function in clinically relevant neoadjuvant and adjuvant settings, both as a monotherapy and in combination with PD-1 blockade, using syngeneic preclinical mouse models.

## Results

### ICOS⁺ Tregs mark an immunosuppressive niche in human breast cancer

To define ICOS expression across immune populations within the human breast tumor microenvironment, we reanalyzed published single-cell transcriptomic data from the integrated Human Breast Cell Atlas (iHBCA) [27], encompassing healthy blood, healthy breast tissue, and treatment-naïve primary breast tumors representing the major subtypes including triple-negative breast cancer (TNBC), estrogen receptor-positive (ER⁺), and HER2-positive (HER2⁺). Across all tissue compartments, ICOS expression was selectively enriched in Tregs relative to other immune cell populations, including CD4⁺ T cells, CD8⁺ T cells, and NK cells (**Fig. 1A**). Within the CD4⁺ T cell compartment, ICOS expression was highest in Tregs compared with T follicular helper (T_FH_, CXCL13⁺), naïve/central memory (CCR7⁺), and naïve/memory (IL7R⁺) subsets (iHBCA annotations; **Fig. 1B**). Within the CD8⁺ T cell compartment, ICOS was most highly expressed on exhausted (LAG3⁺) cells relative to effector (IFNG⁺) and naive/memory (IL7R⁺) subsets (**Fig. 1C**).

**Figure 1.**
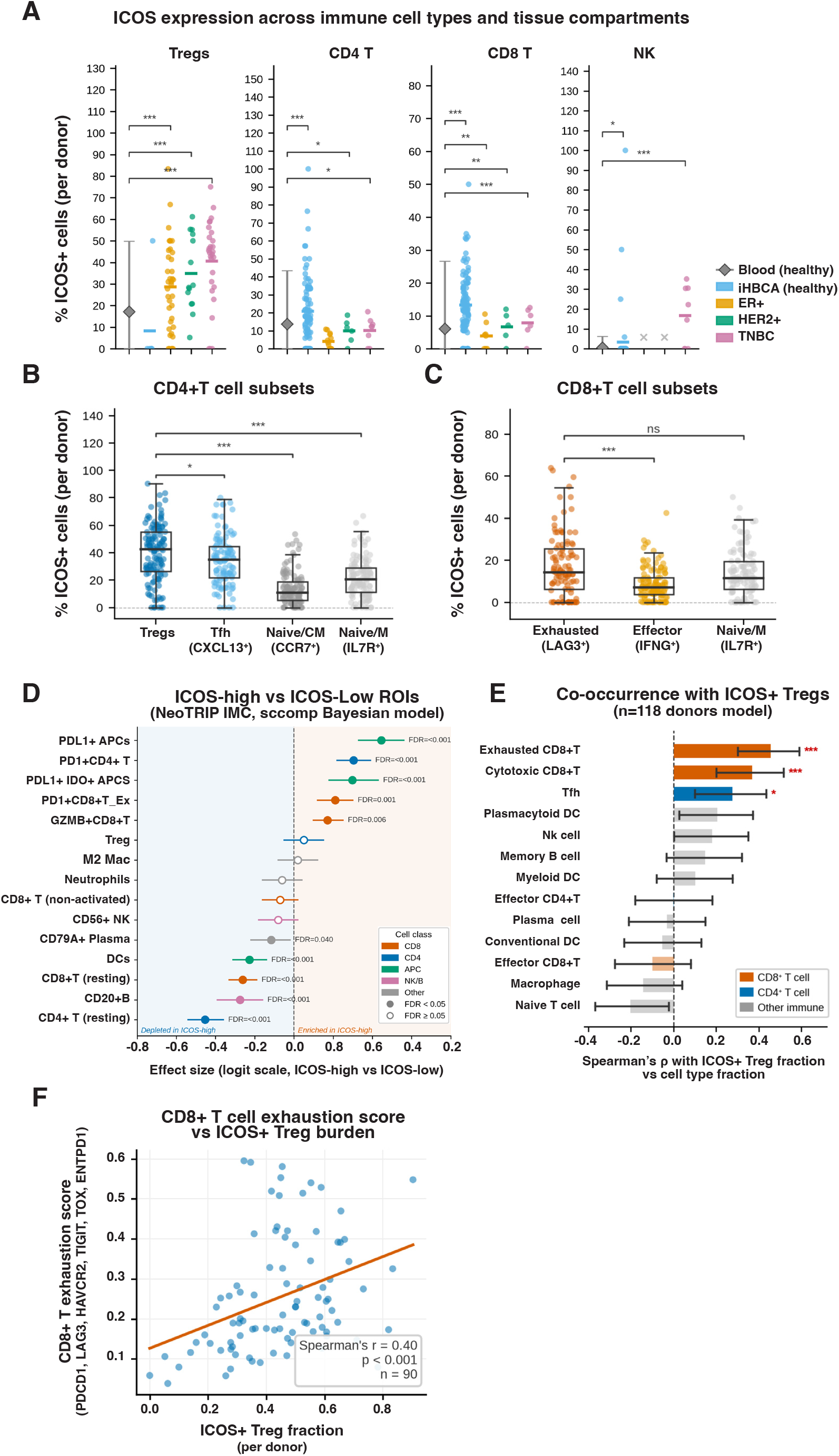
ICOS⁺ Tregs define an immunosuppressive niche in breast cancer. **A.** ICOS expression across major immune cell types (Tregs, CD4⁺ T, CD8⁺ T, NK cells) and tissue compartments. Each dot represents one donor. The y-axis shows % ICOS⁺ cells per donor. Compartments: healthy blood, healthy breast tissue (iHBCA, Reed AD, et al. Nature Genetics 2024), and three breast cancer subtypes (ER⁺, HER2⁺, TNBC) from the integrated Human Breast Cell Atlas (iHBCA). Kruskal-Wallis test with Dunn’s post hoc correction; *, p<0.05; **, p<0.01; ***, p<0.001. **B.** ICOS expression across CD4⁺ T cell subsets, including Tregs, T follicular helper cells (Tfh, CXCL13⁺), naïve/central memory cells (CCR7⁺), and naïve/memory cells (IL7R⁺), in breast cancer donors (iHBCA). Each dot represents one donor. Pairwise Wilcoxon rank-sum test with FDR correction; *, p<0.05; ***, p<0.001. **C.** ICOS expression across CD8⁺ T cell subsets, including exhausted cells (LAG3⁺), effector cells (IFNG⁺), and naive/memory cells (IL7R⁺), in breast cancer donors (iHBCA). Each dot represents one donor. Pairwise Wilcoxon rank-sum test with FDR correction; ***, p<0.001; ns, not significant. **D.** Cell-type abundance differences between ICOS-high and ICOS-low tumor regions of interest (ROIs) in the NeoTRIP IMC dataset (n=1,359 ROIs, Wang, et al. Nature 2023), modeled using Bayesian compositional analysis [29]. Effect sizes are on the logit scale (ICOS-high vs ICOS-low). Points indicate the posterior median, and horizontal bars indicate 95% credible intervals. Dots are colored by cell lineage class (CD8, CD4, APC, NK/B, other). Filled circles, FDR<0.05; open circles, FDR≥0.05. ICOS-high/low was defined by a median split of ICOS expression at the 93rd centile of the global ICOS distribution per ROI. **E.** Co-occurrence of immune cell types with ICOS⁺ Tregs across n=118 donors from the iHBCA, quantified as Spearman’s ρ between ICOS⁺ Treg fraction and each cell-type fraction per donor. Bars are colored by lineage (CD8⁺ T, CD4⁺ T, other immune). Error bars indicate 95% confidence intervals. ***, p<0.001; *, p<0.05 (permutation test, FDR-corrected). **F.** Scatter plot showing the relationship between ICOS⁺ Treg fraction (per donor, n=90) and the CD8⁺ T cell exhaustion score (mean expression of PDCD1, LAG3, HAVCR2, TIGIT, TOX, ENTPD1) in breast cancer donors from the iHBCA. Orange line, linear regression fit. Spearman’s ρ and p-value shown.

Next, we validated ICOS expression at the protein level by multiplex immunohistochemistry performed on an independent cohort of human breast cancer tissues spanning the major subtypes (**Supp. Table 1**). Tumor regions were delineated by hematoxylin and eosin staining (**Supp. Fig. 1A**) and subsequently analyzed for pan-cytokeratin, CD3, CD8, and FoxP3 labeling to score ICOS expression (**Supp. Fig. 1B**). FOXP3⁺ Tregs were preferentially localized to stromal and peri-tumoral niches compared with CD8⁺ T cells in TNBC and ER⁺ tumor tissues, with ICOS⁺ Tregs detected within both the tumor and adjacent stroma regions across all subtypes (**Supp. Fig. 1C-D**). Together, these data identify ICOS as a defining feature of tumor-associated Treg cells in human breast cancer.

We next examined the spatial organization and functional context of ICOS⁺ Tregs within the tumor microenvironment. To this end, we integrated spatial and transcriptional analyses from the NeoTRIP imaging mass cytometry (IMC) dataset [28], comprising 1,359 regions of interest (ROIs) from TNBC patients enrolled in the NeoTRIPaPDL1 trial. Bayesian compositional analysis (sccomp [29]) of ICOS-high versus ICOS-low ROIs revealed significant enrichment of exhausted CD8⁺ T cells and PD-L1⁺ APCs in ICOS-high regions (**Fig. 1D**). ICOS⁺ Tregs exhibited a distinct phenotype relative to ICOS⁻ Tregs, with significantly higher expression of PD-1 (PDCD1) and LAG3 (**Supp. Fig. 2A**). Additionally, ICOS⁺ Tregs expressed significantly higher levels of genes that mark tumor-resident Tregs in human breast cancer and other malignancies [30] [31] [32], including CTLA4, TIGIT, TNFRSF9 (4-1BB), TNFRSF4 (OX40), IL2RA (CD25), CCR8, LAYN, ENTPD1 (CD39), IKZF2 (Helios) and BATF (**Supp. Fig. 2A**). Furthermore, ICOS⁺ Treg abundance was positively correlated with the frequency of PD-L1⁺ APCs (ρ=0.54, p<0.001), IDO1⁺ APCs (ρ=0.47, p<0.001), and PD-L1⁺IDO1⁺ APCs (ρ=0.36, p<0.001) per donor (**Supp. Fig. 2B**), consistent with a co-suppressive microenvironmental niche. Spatial mapping of immune cell centroids confirmed the co-localization of ICOS⁺ Tregs with exhausted CD8⁺ T cells in ICOS-high ROIs, a pattern largely absent in ICOS-low regions (**Supp. Fig. 2C**). Neighbor-graph analysis further demonstrated that ICOS-high Tregs had a significantly lower overall CD8⁺ T cell neighbor fraction in the chemotherapy arm (Δ=-0.016, p=0.010; **Supp. Fig. 2D**), and that the exhausted CD8⁺ T cell share among Treg neighbors was significantly higher in ICOS-high versus ICOS-low ROIs in both the chemotherapy (23.2% vs 18.6%, p=0.010) and chemo-atezolizumab arms (28.6% vs 19.6%, p=0.004) in patients with residual disease (**Supp. Fig. 2E**). To further characterize the relationship between ICOS⁺ Tregs and other immune cell populations, we quantified the co-occurrence of ICOS⁺ Tregs with each immune cell type across n = 118 iHBCA donors using Spearman’s ρ (**Fig. 1E**). ICOS⁺ Treg fraction was positively correlated with CD8⁺ T cell exhaustion score (**Fig. 1F**). Collectively, these data support the model that ICOS⁺ Tregs orchestrate a spatially organized immunosuppressive niche characterized by CD8⁺ T cell exclusion and exhaustion.

**Figure 2.**
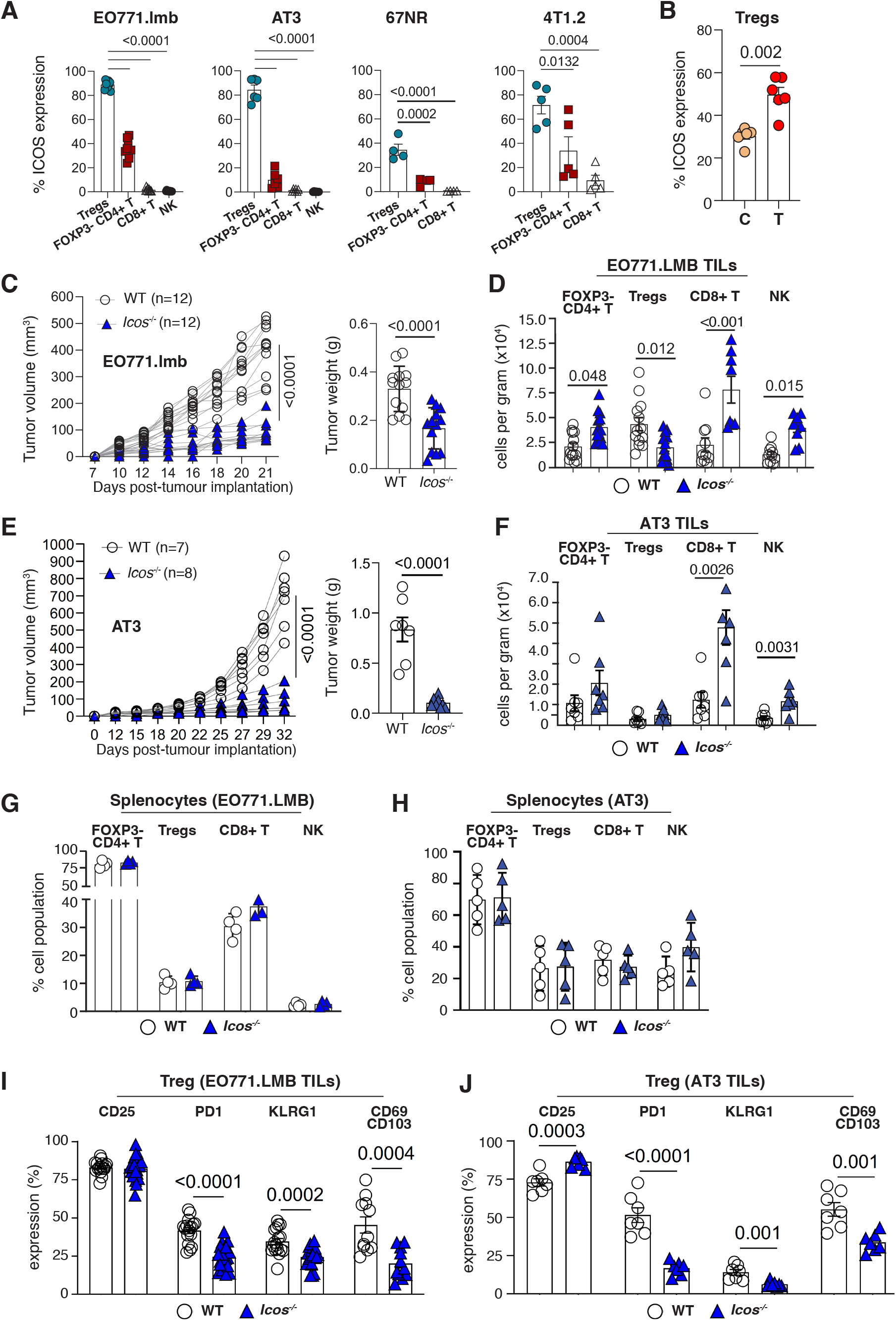
Loss of ICOS alters the mammary tumor microenvironment. **A.** Flow cytometric quantification of ICOS expression across immune subsets in multiple syngeneic murine breast cancer models (EO771.LMB, AT3, 67NR, and 4T1.2), showing ICOS expression in Tregs relative to FOXP3⁻CD4⁺ T cells, CD8⁺ T, and NK cells. Data are presented as mean ± s.e.m. Statistical significance was assessed using ordinary one-way ANOVA (Tukey’s multiple comparison test). p<0.05 was considered significant. **B.** Comparison of ICOS expression on Tregs isolated from healthy contralateral mammary fat pad (C) and EO771.LMB tumor (T). **C** and **E.** Growth kinetics of EO771.LMB and AT3 tumors in wild-type (WT) and *Icos^-/-^* mice were monitored until tumors in WT mice reached approximately 500 mm³ (EO771.LMB) or 900 mm³ (AT3). Quantification of tumor volume and tumor weight is shown in adjacent plots. Mouse numbers, pooled from independent experiments, are shown on the plots. **D** and **F**. Quantification of tumor-infiltrating immune cell populations per gram of tumor tissue, including FOXP3⁻CD4⁺ T, Tregs, CD8⁺ T, and NK cells, in WT and *Icos^-/-^* mice. **G-H.** Flow cytometric analysis of splenocytes isolated from mice analyzed in panels C and E, including FoxP3⁺ Tregs, FoxP3⁻ CD4⁺ T cells, and CD8⁺ T cells expressed as a percentage of TCRβ⁺ cells, and NK cells expressed as a percentage of TCRβ⁻ cells. **I.** Flow cytometric quantification of Treg activation and tissue-residency markers (CD25, PD-1, KLRG1, CD69/CD103) on tumor-infiltrating Tregs from EO771.LMB tumors in WT and *Icos^-/-^* mice. **J.** Flow cytometric quantification of the same Treg markers (CD25, PD-1, KLRG1, CD69/CD103) on tumor-infiltrating Tregs from AT3 tumors in WT and *Icos^-/-^* mice. Data are presented as mean ± s.e.m. Statistical significance was assessed using unpaired two-tailed t-tests. p<0.05 was considered significant.

### ICOS restricts effector immune cell infiltration and promotes mammary tumor growth in mice

To interrogate ICOS activity during mammary tumorigenesis, we employed multiple syngeneic murine mammary tumor models, including EO771.LMB, AT3, 67NR and 4T1.2 [33–36]. Tumors were established by orthotopic implantation of 1×10⁵ tumor cells into the fourth mammary fat pad of 8-12 week-old female C57BL/6 or BALB/c mice and harvested upon reaching volumes of 500 mm³. Consistent with our analyses of human breast tumors, flow cytometric profiling of immune infiltrates within murine mammary tumors (excluding tumor-draining lymph nodes) revealed that ICOS expression within the CD4⁺ T cell compartment was largely confined to Tregs (**Fig. 2A**). In contrast, Tregs isolated from contralateral, non-tumor-bearing mammary glands exhibited relatively lower ICOS expression (**Fig. 2B**), indicating that ICOS upregulation is tumor-context dependent.

To assess the functional relevance of ICOS signaling *in vivo*, we utilized *Icos^-/-^* mice [11]. Systemic loss of ICOS did not impair mammary gland development (**Supp. Fig. 3A**), baseline immune homeostasis, lactation, or nursing capacity, confirming that any tumor-associated phenotypes reflect tumor-specific instead of developmental effects of ICOS deficiency. Both EO771.LMB and AT3 tumors exhibited significantly impaired growth in *Icos^-/-^* mice compared with wild-type C57BL/6 (WT) controls, as demonstrated by reduced tumor volume and lower endpoint tumor weight (**Fig. 2C and E**; **Supp. Fig. 3B and G**). Flow cytometry analysis of EO771.LMB tumors revealed a marked reduction in Treg infiltration (**Fig. 2D**), accompanied by a concomitant increase in effector immune populations, including CD8⁺ T and NK cells (**Fig. 2D**; **Supp. Fig. 3C-D**). Similar expansion of CD8⁺ T and NK cells, accompanied by reduced Treg infiltration, was observed in AT3 tumors from *Icos^-/-^* mice (**Fig. 2F**). In contrast, immune cell composition in the spleen was similar in tumor-bearing WT controls and *Icos^-/-^*mice, indicating that loss of ICOS does not overtly perturb systemic immune homeostasis (**Fig. 2G and H**). Flow cytometric analysis of Treg functional markers revealed that ICOS deficiency reduced PD-1, KLRG1, and CD69/CD103 expression on tumor-infiltrating Tregs in both EO771.LMB and AT3 tumors (**Fig. 2I, J**), indicating impaired Treg activation and loss of tissue residency. In contrast, CD25 expression was unchanged in EO771.LMB tumors and increased in AT3 tumors (**Fig. 2I, J**), suggesting tumor model-dependent regulation of IL-2 signaling. Notably, PD-1, KLRG1 and CD69/CD103 were downregulated in both EO771.LMB and AT3 tumors. This consistency across two distinct tumor models suggests that ICOS selectively maintains the activated, tissue-resident Treg state while leaving IL-2-dependent Treg homeostasis largely intact. Consistent with the expansion of CD8+ T and NK cells in *Icos^-/-^* tumors (**Fig. 2D, F**; **Supp. Fig. 3C, D**), these effector populations exhibited increased proliferation (Ki67) and cytotoxic potential (GzmB) in both EO771.LMB and AT3 tumors (**Supp. Fig. 3E, F, H, I**). Collectively, these findings indicate that ICOS restricts effector immune cell infiltration and promotes mammary tumor growth in mice.

### ICOS remodels the mammary tumor immune landscape, limiting CD8⁺ T cell activation

To define how ICOS signaling shapes the mammary tumor immune landscape, we performed single-cell RNA sequencing (scRNAseq) of CD45⁺ tumor-infiltrating immune cells isolated from EO771.LMB tumors. Immune cells were pooled from three tumors per group derived from either WT controls or *Icos^-/-^*mice. Using established lineage-defining markers, we resolved major immune cell populations within the tumor infiltrate, including Tregs, CD4⁺ and CD8⁺ T cells, B cells, NK cells, dendritic cells, neutrophils, and monocytes (**Fig. 3A**). Comparison of immune cell composition revealed broad remodeling of the tumor immune landscape in the *Icos^-/-^*tumors relative to the WT tumors, indicating that ICOS signaling influences multiple immune compartments, either directly or indirectly. Monocytes and B cell clusters were reduced in *Icos^-/-^* tumors, whereas CD4⁺ T cells were expanded (**Fig. 3B**). In contrast, total CD8+ T, NK, and Treg cell populations exhibited more modest changes in relative abundance. These compositional shifts reflect relative proportions within the pooled scRNAseq dataset. In absolute terms, CD8⁺ T and NK cell numbers quantified by flow cytometry were increased (**Fig. 2D**).

**Figure 3.**
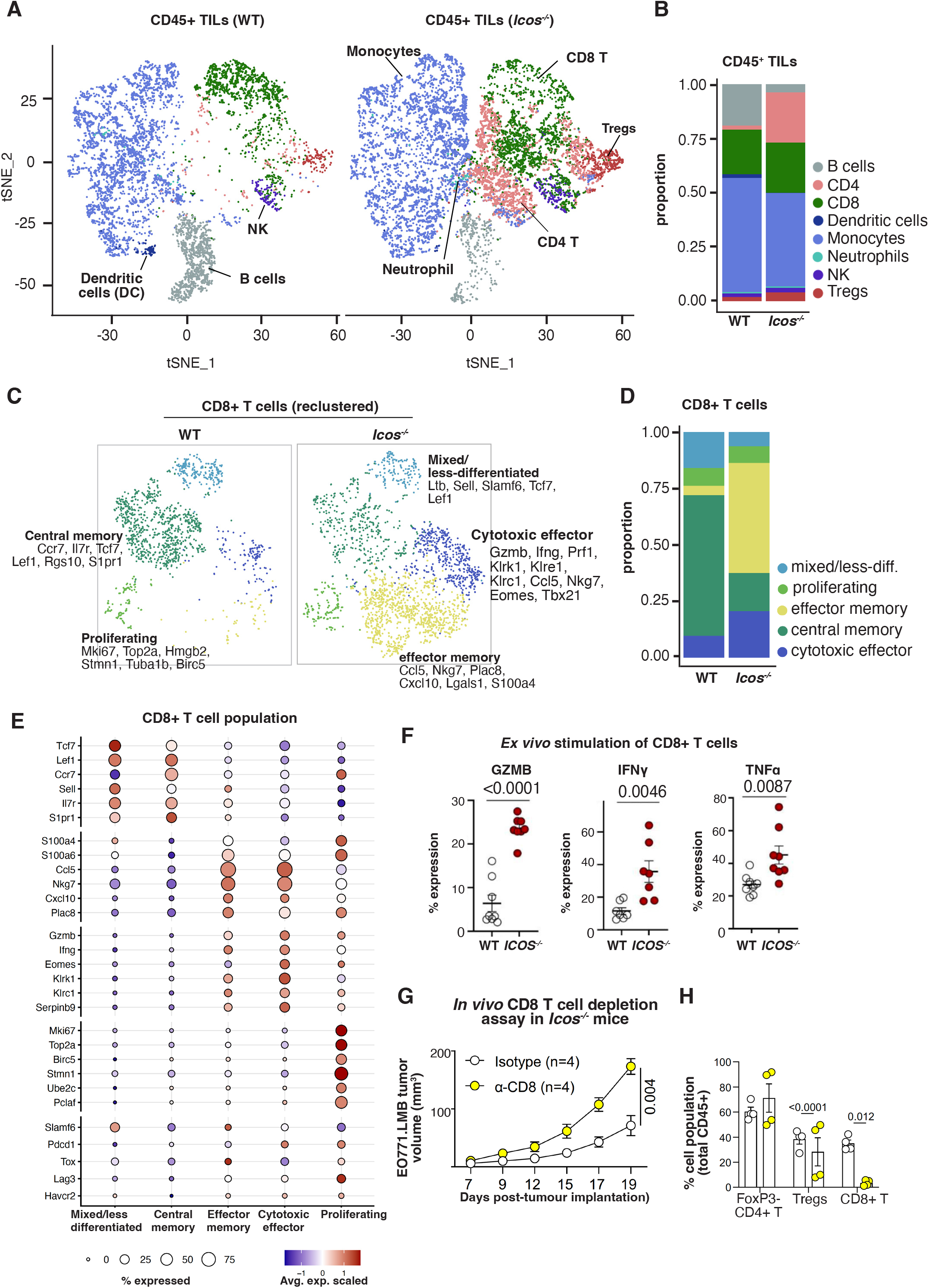
Loss of ICOS reshapes the tumor immune landscape and promotes effector CD8⁺ T-cell differentiation. **A.** t-SNE projections of single-cell RNA sequencing data from tumor-infiltrating CD45⁺ cells isolated from EO771.LMB tumors in wild-type (WT) control and *Icos^-/-^* C57BL/6 mice. Indicated immune cell populations were annotated based on canonical marker gene expression. **B.** Cell type composition within CD45⁺ cells of WT and *Icos^-/-^* mice. **C.** Reclustering of CD8⁺ T cells from the combined dataset reveals distinct transcriptional states, including mixed/less-differentiated (*Ltb, Sell, Slamf6, Tcf7, Lef1*), central memory (*Ccr7, S1pr1, Rgs10*), cytotoxic effector (*Gzmb, Ifng, Prf1, Nkg7*), proliferating (*Mki67, Top2a*), and effector memory (*Ccl5, Plac8, Cxcl10*) subsets, annotated based on top marker genes. **D.** Cell subset composition within CD8⁺ T cells of WT and *Icos^-/-^* mice. **E.** Dot plot of marker gene expression across the five CD8⁺ T cell subsets (combined WT and *Icos^-/-^* dataset), including subset-defining genes and exhaustion-associated genes (*Slamf6, Pdcd1, Tox, Lag3, Havcr2*). Dot size indicates the fraction of expressing cells. Color intensity indicates scaled average expression. **F.** Tumor-infiltrating lymphocytes isolated from WT and *Icos^-/-^* mice were stimulated *ex vivo* with PMA and ionomycin, followed by intracellular cytokine staining. Quantification of granzyme B (GZMB), interferon-gamma (IFNγ), and tumor necrosis factor-alpha (TNFα) in CD8⁺ T cells is shown. **G.** *In vivo* CD8⁺ T cell depletion assay in *Icos^-/-^* mice bearing EO771.LMB tumors. Tumor growth kinetics following isotype or anti-CD8 antibody treatment are shown. **H.** Flow cytometric confirmation of depletion efficiency, showing the frequency of FOXP3⁻CD4⁺ T cells, Tregs, and CD8⁺ T cells among CD45⁺ tumor-infiltrating cells. Data are presented as mean ± s.e.m. Statistical significance was assessed using unpaired two-tailed t-tests. p<0.05 was considered significant.

Marker gene analysis comparing WT and *Icos⁻/⁻* Tregs revealed broad transcriptional differences between the two genotypes **(Supp. Fig. 4)**. As an internal control, *Icos* transcripts were abundant in WT Tregs but depleted in *Icos⁻/⁻* Tregs. Genes reduced in *Icos⁻/⁻* Tregs were dominated by effector and metabolic programs, including the cytotoxic effector molecule *Gzmb*, the glycolytic enzymes *Pkm*, *Eno1* and *Gpi1*, the respiratory chain component *Cox5a*, translational machinery (*Ubb*, *Uba52* and *Eif3f*) and stress-adaptive genes (*Ifitm3* and *Ms4a4b*). Conversely, *Icos⁻/⁻* Tregs showed increased expression of the signaling regulator *Dusp4*, the epigenetic regulator *Hdac7* and the inhibitory molecules *Lag3* and *Nt5e* (CD73). Together, these changes suggest that ICOS helps maintain an activated, metabolically engaged Treg state while restraining selected inhibitory pathways.

Reclustering of CD8⁺ T cells, together with analysis of the top marker genes (**Supp. Fig. 5**), identified five transcriptionally distinct subsets. These comprised a mixed/less-differentiated cluster (*Ltb, Sell, Slamf6, Tcf7, Lef1*), a central memory cluster (*Ccr7, Tcf7, S1pr1, Rgs10*), a cytotoxic effector cluster (*Gzmb, Ifng, Prf1, Nkg7, Eomes, Tbx21*), a proliferating cluster (*Mki67, Top2a, Birc5, Stmn1*), and an effector memory cluster (*Ccl5, Nkg7, Plac8, Cxcl10, S100a4*) (**Fig. 3C-E**; **Supp. Fig. 5**). The mixed/less-differentiated population expressed *Slamf6*, a marker of progenitor-like cells, but showed little or no *Ccr7* expression and lacked prominent cytotoxic or exhaustion-associated gene expression, consistent with a quiescent, less-differentiated state.

Although all CD8⁺ T-cell subsets remained detectable, *Icos*⁻/⁻ tumours showed a marked redistribution of the CD8⁺ compartment. The effector memory (*Ccl5*⁺, *Cxcl10*⁺, low *Gzmb*) and cytotoxic effector (*Gzmb*⁺, *Prf1*⁺) populations were substantially expanded (**Fig. 3C-E**). In contrast, the WT CD8⁺ compartment was dominated by the central memory cluster, which was reduced in *Icos*⁻/⁻ tumours. This shift was accompanied by reciprocal expansion of the effector memory and cytotoxic effector populations, which together became the dominant CD8⁺ T-cell states (**Fig. 3D**). Analysis of exhaustion-associated genes showed that *Pdcd1* and *Tox* were most enriched in the effector memory cluster, with *Pdcd1* also detected in the cytotoxic effector population. *Lag3* expression was largely restricted to the effector memory and proliferating clusters, whereas *Havcr2*, a marker associated with terminal exhaustion, remained low across all subsets (**Fig. 3E**).

To confirm that this transcriptional reprogramming translated into enhanced effector function, we analyzed PMA/ionomycin-stimulated CD8⁺ T cells isolated from WT and *Icos^-/-^* tumors. PMA/ionomycin stimulation bypasses proximal TCR signaling, directly revealing the maximal cytokine-producing capacity acquired by T cells *in vivo* independent of proximal activating cues. CD8⁺ T cells from *Icos^-/-^* tumors exhibited significantly increased intracellular GZMB, TNFα, and IFNγ production (**Fig. 3F**). Furthermore, *in vivo* antibody-mediated depletion of CD8⁺ T cells in EO771.LMB tumor-bearing *Icos^-/-^*mice abrogated the tumor-suppressive effect associated with ICOS loss, leading to accelerated tumor growth comparable to control conditions (**Fig. 3G**). This treatment achieved efficient and selective depletion of the targeted CD8⁺ T cell population within the tumor (**Fig. 3H**). Overall, these findings link the transcriptional remodeling to functional anti-tumor immunity dependent on cytotoxic CD8⁺ T cells.

### Anti-ICOS therapy remodels the TME and suppresses primary tumor growth

We next tested the therapeutic efficacy of ICOS blockade using a monoclonal antibody (mAb, clone 7E.17G9) in WT C57BL/6 female mice bearing EO771.LMB mammary tumors (**Fig. 4A**). Mice were randomized into two groups once tumors became palpable (∼day 9-10) and received either 10 mg/kg of anti-ICOS mAb or isotype control via intraperitoneal injection every three days for a total of three doses. Tumors were harvested once the majority of isotype control tumors reached the experimental endpoint (∼500 mm³). As an additional control, we included untreated *Icos^-/-^*tumor-bearing mice to benchmark the effects of genetic versus pharmacological ICOS inhibition.

**Figure 4.**
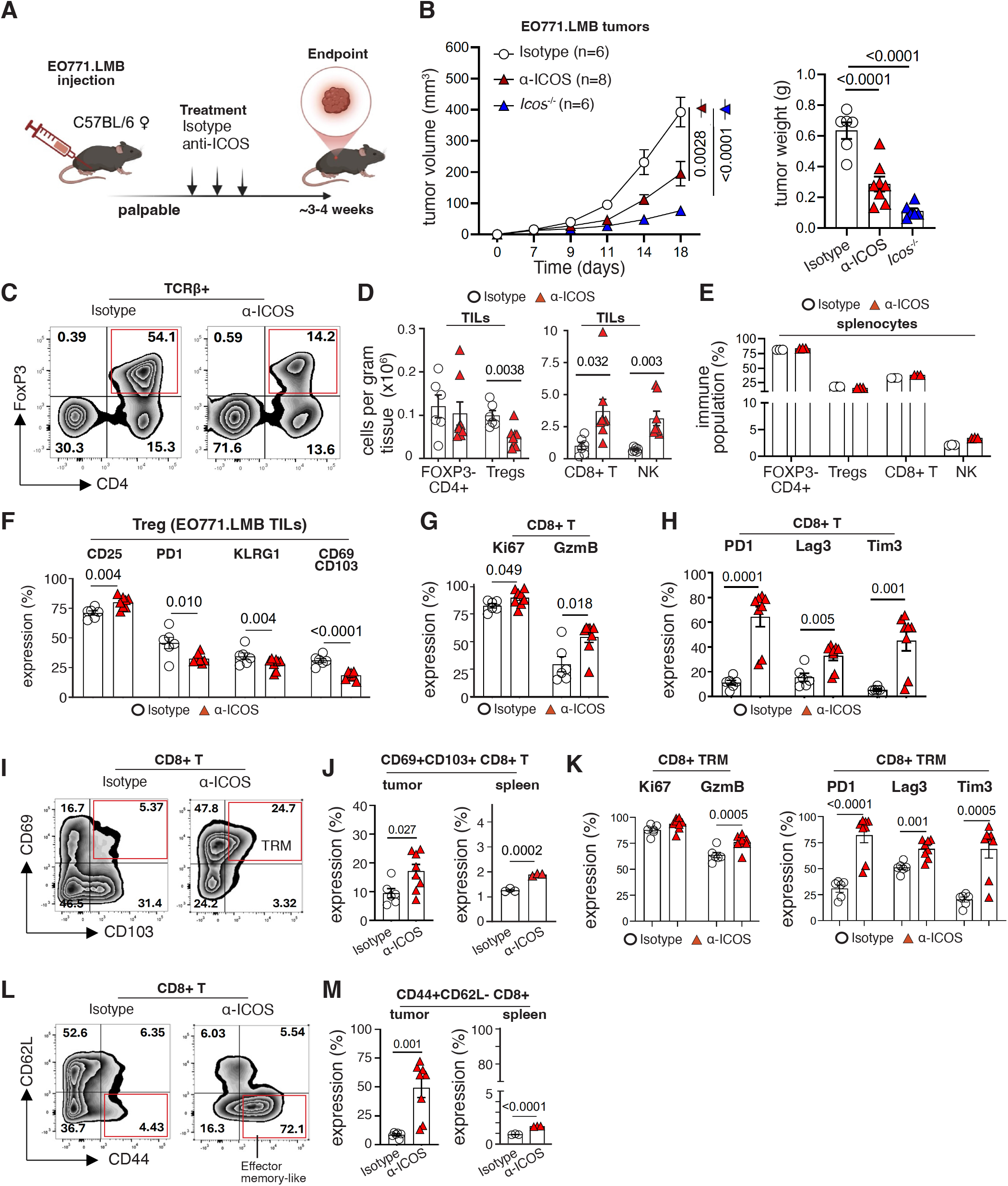
Pharmacological blockade of ICOS enhances anti-tumor immunity in breast cancer. **A.** Schematic of the treatment regimen. **B.** Tumor growth kinetics and endpoint tumor weight in EO771.LMB tumor-bearing wild-type (WT) C57BL/6 mice treated with anti-ICOS monoclonal antibody (clone 7E.17G9, 10 mg/kg, intraperitoneally, every 3 days for three doses) or isotype control. Untreated *Icos^-/-^* tumor-bearing mice were included as a genetic reference control. **C.** Representative flow cytometry plots of FoxP3+ Tregs. **D.** Flow cytometric quantification of tumor-infiltrating lymphocytes (TILs), including FoxP3+ Tregs, FoxP3-CD4+ conventional T cells, CD8+ T cells, and NK cells, following anti-ICOS or control treatment. **E.** Flow cytometric analysis of splenocyte immune populations (FoxP3+ Tregs, FoxP3-CD4+ T cells, CD8+ T cells, and NK cells) following anti-ICOS or control treatment. FoxP3⁺ Tregs, FoxP3⁻ CD4⁺ T cells, and CD8⁺ T cells are presented as a percentage of TCRβ⁺ cells, and NK cells as a percentage of TCRβ⁻ cells. **F.** Flow cytometric analysis of intratumoral Treg functional markers (CD25, PD-1, KLRG1, CD69/CD103) following anti-ICOS treatment. **G.** Flow cytometric analysis of Ki67 and GzmB expression in CD8+ T cells. **H.** Flow cytometric analysis of PD-1, Lag3, and Tim3 expression in CD8+ T cells. **I.** Representative flow cytometry plots of CD69 and CD103 expression on CD8+ T cells. **J.** Quantification of CD69+CD103+ tissue-resident memory (TRM) CD8+ T cells in tumor and spleen. **K.** Flow cytometric analysis of Ki67, GzmB, PD-1, Lag3, and Tim3 expression in CD8+ TRM cells. **L.** Representative flow cytometry plots of CD44 and CD62L expression on CD8+ T cells. **M.** Quantification of CD44+CD62L-effector-memory CD8+ T cells in tumor and spleen. Data are presented as mean +/-s.e.m. Statistical significance was determined using unpaired two-tailed t-tests. p<0.05 was considered significant.

Anti-ICOS treatment significantly reduced tumor growth and final tumor weight compared to isotype control (**Fig. 4B**; **Supp. Fig. 6A**), with outcomes similar to those observed in *Icos^-/-^* mice, indicating that transient ICOS blockade is sufficient to restrict tumor progression. Anti-ICOS treatment also reduced surface ICOS expression on intratumoral FOXP3⁻CD4⁺ T cells, Tregs and CD8⁺ T cells (**Supp. Fig. 6B**), confirming on-target antibody activity. Flow cytometry analysis revealed that anti-ICOS treatment led to a reduction in the FoxP3⁺ Treg population, accompanied by increased infiltration of CD8⁺ T cells and NK cells (**Fig. 4C-D**), mirroring effects seen with EO771.LMB tumors isolated from *Icos^-/-^*female mice (**Fig. 2D**). Importantly, Treg, CD8⁺ T and NK cell frequencies in the spleen were largely unchanged (**Fig. 4E**), indicating that the treatment exerted tumor-specific compositional changes. Furthermore, anti-ICOS treatment attenuated the intratumoral Treg activation phenotype, with decreased PD-1 and KLRG1 expression and loss of the tissue-residency markers CD69 and CD103 (**Fig. 4F**), as well as reduced frequencies of GzmB⁺ Tregs (**Supp. Fig. 6C**). KLRG1 marks highly activated, suppressive Treg subsets within tumors [37] [38], supporting the interpretation that ICOS blockade dismantles the activated Treg state. In contrast, anti-ICOS treatment did not reduce CD25 expression on tumor-infiltrating Tregs (**Fig. 4F**).

Analysis of cytotoxic lymphocytes revealed increased CD8⁺ T cell activation following ICOS blockade. This reprogramming was associated with increased expression of Ki67, GzmB and the exhaustion markers PD-1, Lag3 and Tim3 in the CD8⁺ T cell compartment (**Fig. 4G-H**), consistent with the effector differentiation observed in *Icos^-/-^* tumors (**Fig. 3C-E**). Furthermore, the ICOS blockade resulted in enrichment of CD69⁺CD103⁺ tissue-resident memory (TRM) CD8⁺ T cells in primary tumor site and a modest but significant increase in spleen (**Fig. 4I-J**). These cells showed increased expression of GzmB and exhaustion markers in TILs (**Fig. 4K**), consistent with heightened local activation. Furthermore, ICOS inhibition drove a pronounced shift in CD8⁺ T-cell differentiation, characterized by enrichment of CD44^+^CD62L^-^ effector-memory cells in TILs (**Fig. 4L-M**). Together, these data demonstrate that pharmacological ICOS blockade phenocopies genetic ICOS loss, dismantling the intratumoral Treg niche while licensing cytotoxic, effector-memory and tissue-resident memory CD8⁺ T cell responses that confer durable systemic anti-tumor immunity.

### ICOS blockade alone improves the overall survival rate in metastatic settings

Metastatic disease responds poorly to treatment and is a major cause of breast cancer mortality. Thus, in addition to controlling primary tumor growth, an important therapeutic objective is to treat and limit metastatic spread. Using the EO771.LMB orthotopic mammary tumor model, which reliably develops spontaneous lung metastases 7-8 weeks after surgical resection, we investigated the impact of ICOS blockade on metastatic progression (**Fig. 5A, B**). Flow cytometry of lung-infiltrating immune cells revealed that metastatic lesions were characterized by a marked enrichment of Tregs, especially the ICOS⁺ subset (**Fig. 5C, D**), and a concomitant reduction in CD8⁺ T cells (**Fig. 5E**; **left panel**). Moreover, CD8⁺ T cells in metastatic lung tissues expressed elevated levels of PD-1 (**Fig. 5E**; **right panel**), consistent with an exhausted phenotype and an immunosuppressive metastatic microenvironment.

**Figure 5.**
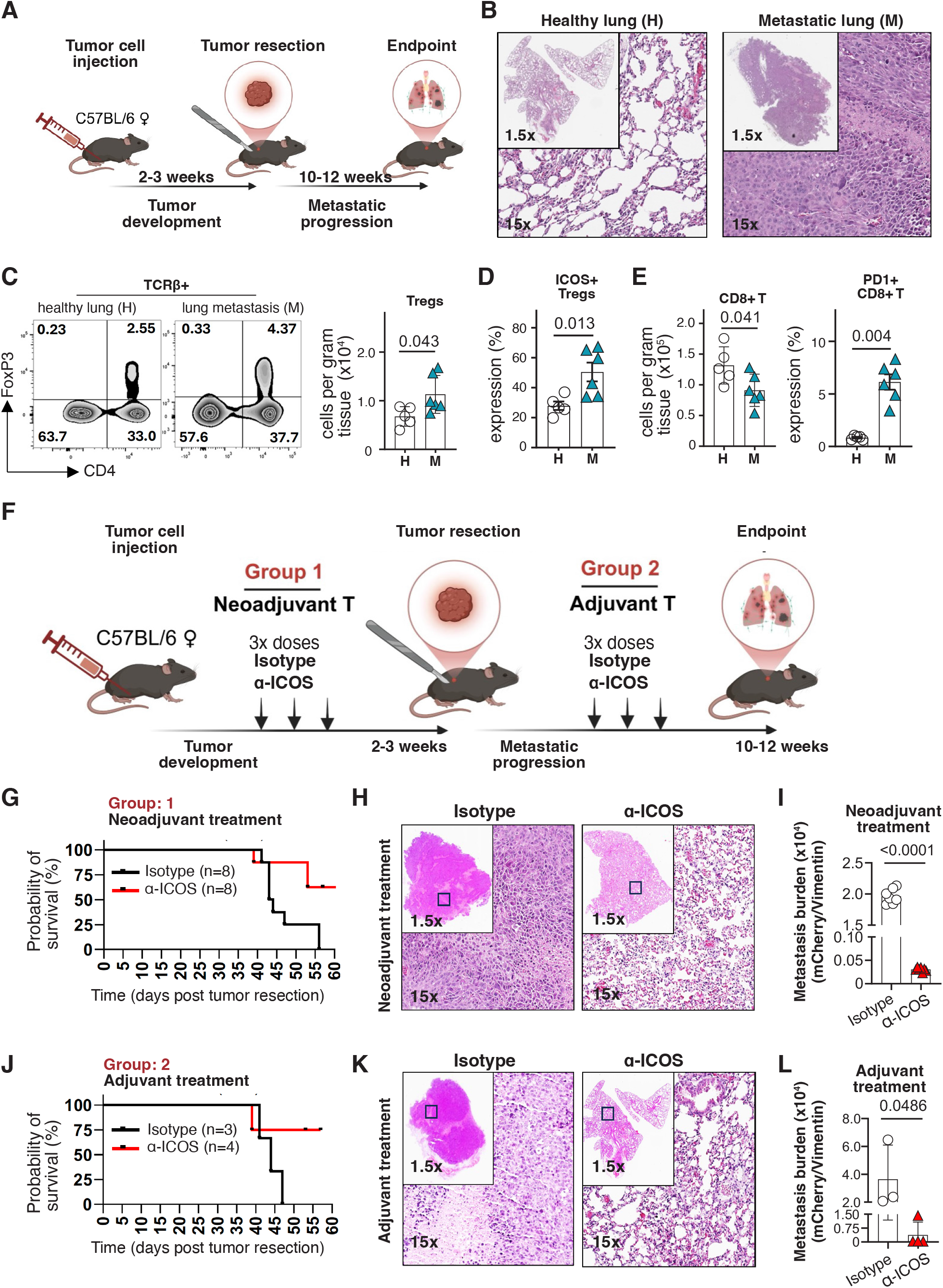
ICOS blockade suppresses spontaneous lung metastasis and improves survival in neoadjuvant and adjuvant breast cancer settings. A. Schematic of the EO771.LMB spontaneous metastasis model. B. Representative images of healthy and metastatic lung tissue. C-D. Flow cytometric analysis of Tregs and ICOS expression on Tregs isolated from lungs of healthy and tumor-bearing mice. E. Quantification of CD8+ T cells and PD-1 expression on CD8+ T cells in the metastatic lung microenvironment. F. Schematic of neoadjuvant and adjuvant treatment strategies. For neoadjuvant therapy, EO771.LMB tumor-bearing wild-type C57BL/6 mice received three doses of anti-ICOS monoclonal antibody (10 mg/kg) or isotype control prior to surgical resection of the primary tumor. For adjuvant therapy, the same dosing regimen was administered following tumor resection. G and J. Kaplan-Meier survival analysis of mice randomized to receive isotype control or anti-ICOS treatment, evaluating therapeutic efficacy in both neoadjuvant and adjuvant treatment settings. H and K. Representative H&E-stained lung sections showing metastatic burden across treatment groups. I and L. Quantitative PCR analysis detecting tumor-derived mCherry transcripts in lung tissues as an independent measure of metastatic burden. Data are presented as mean +/-s.e.m. Statistical significance was assessed using unpaired two-tailed t-tests unless otherwise indicated. p<0.05 was considered significant.

We designed two clinically relevant intervention strategies involving neoadjuvant (pre-surgical) and adjuvant (post-surgical) administration of anti-ICOS monoclonal antibody (mAb) (**Fig. 5F**). In the neoadjuvant setting, WT mice bearing palpable EO771.LMB tumors received three doses of Isotype or anti-ICOS mAb before surgical resection, whereas in the adjuvant arm, the same dosing regimens were administered one-week post-resection of tumor. Kaplan-Meier survival analysis showed that anti-ICOS monotherapy improved survival compared to isotype controls in both neoadjuvant and adjuvant therapies (**Fig. 5G and J**). Histological assessment (**Fig. 5H and K**) and mCherry-specific qPCR (**Fig. 5I and L**) revealed significantly fewer and smaller metastatic foci in both neoadjuvant-and adjuvant-treated mice compared to isotype controls. These data demonstrate that ICOS blockade, administered before or after surgical resection, durably suppresses metastatic outgrowth and extends survival.

To assess whether ICOS blockade augments immune checkpoint inhibition, EO771.LMB tumor-bearing mice were treated with anti-ICOS, anti-PD-1, or the combination. Notably, dual anti-ICOS and anti-PD-1 treatment resulted in improved tumor control, with reduced tumor volume and endpoint tumor weight relative to either single agent (**Fig. 6A**; **Supp. Fig. 6D, E**). To determine whether these findings extended beyond a single model, we evaluated anti-ICOS and anti-PD-1 combination therapy in the 4T1.2 model. The single and combination treatments showed similar primary tumor control as in EO771.LMB tumors (**Fig. 6B**; **Supp. Fig. 6H, I**).

**Figure 6.**
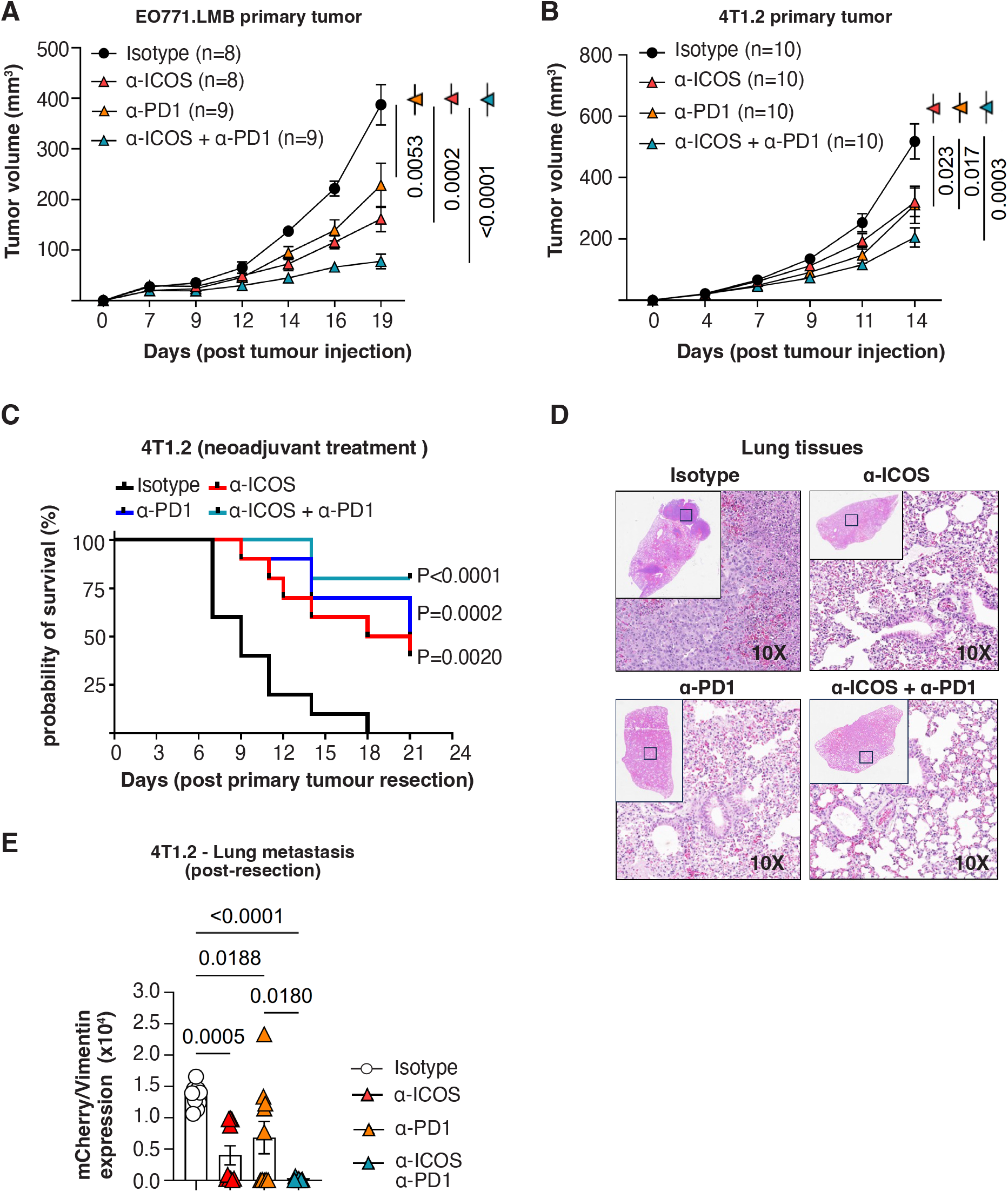
Blockade of ICOS and PD-1 shows comparable effects in a neoadjuvant breast cancer model. **A.** EO771.LMB mammary tumors were established in female C57BL/6 mice. Once tumors were palpable, mice were treated intraperitoneally with anti-ICOS (10µg/g), anti-PD-1 (10µg/g), the combination of both antibodies, or isotype control antibodies, administered every three days for a maximum of three doses. Tumor volumes were monitored longitudinally. **B.** 4T1.2 mammary tumors were established in female BALB/c mice and treated with the same regimen; tumor volumes were monitored longitudinally. **C.** Kaplan-Meier survival analysis of 4T1.2 tumor-bearing mice in the neoadjuvant setting, comparing isotype control, anti-ICOS, anti-PD-1, and the combination. **D.** Representative hematoxylin and eosin (H&E) staining of lungs showing metastatic burden across 4T1.2 treatment groups. **E.** Quantification of lung metastatic burden by qPCR estimation of mCherry expression by 4T1.2 cells across treatment groups. Data are presented as mean +/-s.e.m. Statistical significance was determined using unpaired two-tailed t-tests (tumor growth and metastasis quantification) and the log-rank test (survival analysis). p<0.05 was considered significant.

Beyond primary tumor control, ICOS blockade and the combination reshaped the intratumoral effector CD8⁺ T cell compartment in both models. In EO771.LMB tumors, anti-ICOS increased the frequencies of PD-1⁺ and Ki67⁺ CD8⁺ T cells, while the combination drove the strongest expansion of GzmB⁺ and CD44⁺CD62L⁻ effector-memory CD8⁺ T cells **(Supp. Fig. 6F, G)**. Similarly, in 4T1.2 tumors, anti-ICOS increased PD-1⁺ CD8⁺ T cells **(Supp. Fig. 6J)**, and all treatment arms elevated GzmB⁺ CD8⁺ T cells, with the combination showing the strongest induction and also significantly increasing CD8⁺ T cell proliferation **(Ki67; Supp. Fig. 6J)**.

Following resection of 4T1.2 primary tumors, Kaplan-Meier survival analysis showed that anti-ICOS and anti-PD-1 monotherapies, as well as their combination, significantly improved survival compared with isotype controls in the neoadjuvant setting (**Fig. 6C**). Histological assessment revealed significantly fewer metastatic foci in neoadjuvant-treated mice compared with isotype controls (**Fig. 6D**), consistent with the reduced mCherry signal from 4T1.2 cells detected by qPCR (**Fig. 6E**). Together, these data show that ICOS blockade, alone or in combination with PD-1 blockade, relieves Treg-mediated suppression and enhances CD8⁺ T cell-mediated tumor control.

## Discussion

An immunosuppressive tumor microenvironment (TME) limits the effectiveness of immune checkpoint inhibitors in breast cancer. Our study shows that ICOS signaling is not simply a marker of T cell activation. It is required to sustain an immunologically cold TME. Disrupting this axis, whether through genetic ablation or pharmacological blockade, remodels the TME into a more immune-permissive state, marked by stronger cytotoxic activity among CD8⁺ T cells.

We found that ICOS expression is selectively enriched on tumor-infiltrating FOXP3⁺ Tregs across several breast cancer subtypes. Intra-tumoral Tregs expressed significantly more ICOS than Tregs from healthy mammary tissue or peripheral blood. In human breast tumors, CTLA4, TIGIT, TNFRSF9/4-1BB and ENTPD1/CD39 were all more highly expressed in ICOS⁺ Tregs (**Supp. Fig. 2A**). In line with this, tumor-infiltrating Tregs from *Icos^-/-^* EO771.LMB murine tumors showed reduced expression of genes tied to the activated Treg state (**Supp. Fig. 4**). These included *Gzmb*, the suppressive effector molecule that tumor-associated Tregs use to eliminate NK and CD8⁺ T cells [39]; the glycolytic enzymes *Pkm*, *Eno1* and *Gpi1*, which meet the heightened bioenergetic demands of activated Tregs; and the activation markers PD-1 and KLRG1 (**Fig. 2I, J**). By contrast, CD25 expression on Tregs was maintained or even increased (**Fig. 2I, J and Fig. 4F**). This suggests that IL-2-dependent homeostatic signaling remains intact even as ICOS is lost. Furthermore, ICOS loss also reduced the fraction of Tregs co-expressing the tissue-retention markers CD69 and CD103 [40] [41]. Collectively, these markers characterize highly activated, suppressive tumor-infiltrating Tregs and were downregulated across both genetic and pharmacological disruption of ICOS, indicating that ICOS selectively maintains the activated Treg state [37] [38] [19]. The loss of these markers likely reflects a shift away from the suppressive Treg state rather than impaired Treg survival. This fits with mechanistic studies showing that ICOS signaling is needed to sustain FOXP3-driven transcriptional programs and suppressive function in Tregs [12–17].

Beyond the Tregs themselves, this suppressive niche has a defined spatial structure in human breast tumors. The NeoTRIP imaging mass cytometry cohort included 1,359 tumor regions from 279 patients [28]. Within this cohort, ICOS-high tumor regions were enriched for exhausted CD8⁺ T cells as well as for PD-L1⁺ and IDO1⁺ antigen-presenting cells (**Fig. 1D**). At the donor level, ICOS⁺ Treg abundance correlated with the frequencies of PD-L1⁺, IDO1⁺ and PD-L1⁺IDO1⁺ antigen-presenting cells (n=118 donors, **Supp. Fig. 2B**). It also co-occurred most strongly with exhausted and cytotoxic CD8⁺ T cells across iHBCA donors (**Fig. 1E**) and tracked with the CD8⁺ T cell exhaustion score (**Fig. 1F**). This niche was linked to treatment outcome as well. ICOS⁺ Tregs had fewer CD8⁺ T cell neighbors in the chemotherapy arm (**Supp. Fig. 2D**), a pattern that held among patients with residual disease but not among those with a pathological complete response. CD8⁺ T cells neighboring ICOS⁺ Tregs were more frequently exhausted in both the chemotherapy and chemo-atezolizumab arms (**Supp. Fig. 2E**). Together, these findings place ICOS⁺ Tregs within the exhausted, PD-L1-high microenvironment that immune checkpoint blockade targets and link this niche to incomplete response to neoadjuvant therapy. These human spatial analyses are correlative and will require validation in independent patient cohorts.

Given that clinically relevant ICOS-targeting antibodies act systemically, we used systemic ICOS-deficient mice rather than Treg-specific deletion models. This approach captures the combined effects of ICOS inhibition across the tumor immune ecosystem. Because it perturbs ICOS in all lineages at once, though, lineage-restricted deletion will still be needed to resolve the relative contributions of the Treg, CD8⁺ and CD4 compartments. Our findings are consistent with previous studies showing that systemic ICOS loss or Treg-restricted ICOS deficiency enhances antitumour immunity by disrupting ICOS-dependent regulatory interactions between Tregs and cytotoxic lymphocytes. [21–24, 42]. Together, this implicates Treg-intrinsic ICOS signaling as the main driver of immune suppression.

A recent study identified ICOS as a key regulator of exhausted CD8⁺ T cells. It restrains their memory-like properties and functional capacity, both during chronic viral infection and in a mouse model of hepatocellular carcinoma [43]. In the absence of ICOS, we observed a marked expansion of two CD8⁺ T cell populations at the expense of central-memory cells. These were cytotoxic-effector cells, expressing Gzmb, Ifng, Prf1, Nkg7, Eomes and Tbx21, and effector-memory cells, expressing Ccl5, Cxcl10, Nkg7, Plac8 and S100a4. This shift was accompanied by increased production of granzyme B, TNFα and IFNγ upon *in vitro* restimulation (**Fig. 3C-F**). These findings suggest that ICOS restricts effector differentiation and cytotoxic capacity within the breast tumor immune microenvironment. Pdcd1 and Tox were most enriched in the expanded effector-memory cluster, while Lag3 was enriched in both the effector-memory and proliferating clusters (**Fig. 3E**). In the human breast tumor microenvironment, ICOS expression was highest in LAG3⁺ exhausted CD8⁺ T cells (**Fig. 1C**). This raises the possibility that ICOS blockade enhances antitumor immunity through two routes, by disrupting ICOS⁺ Tregs and by modulating exhausted CD8⁺ T cells directly. ICOS blockade also enriched CD8⁺ tissue-resident memory (TRM) cells, in association with improved tumor control. This is particularly relevant in breast cancer, where greater intratumoral TRM-cell abundance has been linked to favorable clinical outcomes [44–46].

One notable translational feature of our study is that ICOS blockade worked as a single agent. Anti-ICOS monotherapy significantly restricted primary mammary tumor growth, to a degree comparable with genetic Icos deletion. In the metastatic setting, it also reduced spontaneous lung metastatic burden and significantly prolonged survival, whether given as a neoadjuvant or adjuvant regimen. Since metastatic disease is the leading cause of death in breast cancer, activity against both primary and metastatic disease as a single agent is a clinically meaningful feature of this approach.

Our findings build on previous work with ICOS-targeting antibodies in cancer. KY1044 is an effector-competent IgG1 antibody that depletes ICOS-high Tregs. It showed antitumor activity and improved the efficacy of anti-PD-L1 therapy in subcutaneous syngeneic tumor models, including the colorectal carcinoma models CT26.WT and MC38. This combination, however, did not work against subcutaneous B16-F10 melanoma or 4T1 mammary tumors [47]. In a humanized mouse model of breast cancer, a neutralizing anti-human ICOS antibody reduced Treg numbers but controlled tumor growth only when combined with cyclophosphamide [48]. Another study found that anti-ICOS blockade worked in an MLL3-mutant mouse mammary tumor model, depleting tumor-infiltrating Tregs and inhibiting primary tumor development [49]. Notably, earlier studies did not examine orthotopic breast tumor models of spontaneous metastasis, and none assessed survival following neoadjuvant or adjuvant ICOS blockade. Our study addresses these gaps and establishes CD8⁺ T cells as the required effectors of anti-ICOS therapy. As with these antibodies, ours may act at least in part by depleting ICOS-high Tregs rather than purely by blocking ICOS signaling. ICOS inhibition, though, did not cause overt systemic immune dysregulation, which suggests a favorable therapeutic index in both neoadjuvant and adjuvant settings.

In breast cancer, the limited clinical benefit of PD-1 monotherapy points to a need for rational combination strategies, and our data suggest that combining ICOS and PD-1 blockade may offer additive benefit. PD-1 inhibition reinvigorates exhausted effector lymphocytes, while ICOS blockade removes the dominant Treg barrier that limits effector differentiation and cytotoxic activity in tumor-infiltrating T cells. Dual blockade trended toward better tumor control than either monotherapy alone, though the benefit was modest and will need further characterization in larger preclinical and clinical studies.

In summary, our findings support the clinical development of ICOS-targeted strategies as a way to sensitize immunologically cold breast tumors to immune checkpoint blockade and improve long-term outcomes across breast cancer subtypes. ICOS-marked effector Tregs have been reported in several cancers [21, 22, 25, 26]. Our study extends this concept to breast cancer. We define the spatial relationship between ICOS⁺ Tregs and exhausted CD8⁺ T cells, show single-agent efficacy of ICOS blockade against orthotopic primary tumors and spontaneous metastases, and demonstrate a survival benefit in both neoadjuvant and adjuvant regimens. The efficacy of ICOS blockade across two tumor models, two genetic backgrounds (C57BL/6 and BALB/c) and a range of immunological contexts supports the generalizability of this therapeutic strategy.

## Author contributions

Conceptualization: C.L., S.G., A.V., S.M., B.P.

Investigation: C.L, S.G., W.J.H., L.N, K.T, Y.G.

Writing-original draft: S.M., B.P.

Writing-review & editing: S.V., V.Y., R.L.A., D.M., L.A.M., J.M., C.L., C.K., L.M., B.Y., A.V., A.K., S.M., B.P.

## Acknowledgement

We thank Rebecca Brown for assistance with optimisation of the immunohistochemistry antibody panel, David Baylon for support with flow cytometry workflows, and our patient advocates, Jane Miller and Ian Sadler, for their valuable feedback.

## Material and methods

### Human samples

Human breast tissues were obtained from consenting patients through the Austin Hospital with relevant institutional review board approval. Human Ethics approval was obtained from the Austin Health Human Research Ethics Committee (HREC/66494/Austin-2020). The sample information, is provided in **Supplementary Table 1**.

### Cell culture

The primary cell line used in this study was EO771.LMB, a highly metastatic variant derived from a lung nodule of a C57BL/6 mouse bearing an EO771 tumor [34]. In addition, AT3 (a metastatic mammary carcinoma line derived from C57BL/6 mice), 4T.1 (highly metastatic mammary carcinoma line derived from BALB/c mice) and 67NR (a non-metastatic mammary carcinoma line derived from BALB/c mice) were also used for comparative studies. All cell lines were cultured in Dulbecco’s Modified Eagle’s Medium (DMEM, Gibco™, Cat no: 11965092) supplemented with 10% foetal bovine serum (FBS, Bovogen, Cat no: SFBS), 10 mM HEPES (Gibco™, Cat no: 15630130), and 1% penicillin/streptomycin (Pen Strep, Gibco™, Cat no: 15140122), and maintained in a humidified incubator at 37 °C with 5% CO₂. To ensure consistency in intra-mammary fat pad injections, cells were subjected to two rounds of passaging before injecting them *in vivo*. For long-term storage, cells were cryopreserved in freezing medium containing 10% dimethyl sulfoxide (DMSO, EMD Millipore, Cat no: 317275) and 90% FBS, initially stored at -80 °C and subsequently transferred to liquid nitrogen.

### Cell preparation for intra-mammary fat pad injections

For intra-mammary fat pad injections, cells were harvested at 70-80% confluency using TrypLE™ Express Enzyme (1X) (Gibco™, Cat no: 12604021), washed with sterile Phosphate Buffered Saline (PBS, Gibco™, Cat no: 10010023), and pelleted by centrifugation at 500g for 5 min at 4°C. The pellet was resuspended in PBS, and viability was assessed by Trypan Blue (Gibco™, Cat no: 15250061) exclusion using a TC10™ Automated Cell Counter (BioRad), with 80-90% viability required for injections. Cell suspensions were adjusted to ∼10×10⁶ cells/mL in PBS, corresponding to ∼1×10⁵ cells per 10 µL injection.

### Animal studies

All animal procedures were approved by the Austin Health Animal Ethics Committee (AEC A2021-05727, A2022-05785, A2023-05824, Melbourne) and conducted in accordance with the Australian Code of Practice for the Care and Use of Animals for Scientific Purposes. Mice were housed under specific pathogen-free conditions in a temperature-controlled facility with ad libitum access to food and water. *Icos^-/-^*mice [11] were obtained from Dr Zhenjun Chen (Peter Doherty Institute, Melbourne).

Orthotopic tumor growth and metastasis experiments were performed as previously described. Briefly, EO771.LMB and AT3 murine breast cancer cells were injected into the fourth mammary fat pad of female C57BL/6 mice, and 4T1.2 and 67NR cells were implanted into female BALB/c mice, to establish orthotopic mammary tumors. Tumor growth was monitored using digital calliper, and tumor volume was calculated using the formula ½ × (length × width²). At experimental or ethical endpoints, primary mammary tumors were excised and weighed. For metastatic dissemination studies, primary tumors were surgically resected once they reached a volume of 250-500 mm³. Mice were monitored until the experimental endpoint, at which time lungs were harvested for assessment of metastatic burden. All five lobes of the lung tissues were homogenised in 1 ml of TBST (1× Tris-buffered saline with 0.5% (vol/vol) Triton X-100). Genomic DNA was extracted using the AllPrep DNA/RNA Mini Kit (Qiagen; Cat no: 80204) according to the manufacturer’s instructions. Metastatic burden was quantified by duplex quantitative PCR detection of tumor cell-specific mCherry gene (FAM-conjugated probe) and a housekeeping vimentin gene (VIC-conjugated). The qPCR reaction was set up with 20ng DNA template, 1X Universal TaqMan Mix, 0.05 µM Probe mix in 10 µL reaction mix, and the relative tumor burden was quantified as 10000 x 2^-ΔCT^.

### In vivo antibody treatment

In vivo antibody treatment commenced one week after tumor cell inoculation, corresponding to the early stage of tumor growth. Mice received intraperitoneal (IP) injections of isotype control antibody (InVivoMAb rat IgG2a anti-trinitrophenol; Bio X Cell, Cat no: BE0089), anti-mouse ICOS (InVivoMAb; Bio X Cell, Cat no: BE0059), anti-mouse PD-1 (CD279) (InVivoMAb; Bio X Cell, Cat no: BE0273), anti-mouse NK1.1 (InVivoMAb; Bio X Cell, Cat no: BE0036), or anti-mouse CD8α (InVivoMAb; Bio X Cell, Cat no: BE0117). Antibodies were administered at 10 µg/g body weight in a maximum injection volume of 100 µL.

Treatments were delivered every third day for a maximum of three doses. For combination treatment groups, the two antibodies were given as separate IP injections one hour apart.

### Cell preparation for intracellular staining

Expression of various immune cell markers (**Supp. Table 2**) was measured using standard flow cytometry. The cells isolated from digested tissue were transferred to a 96-well plate and spun down at 1500 rpm for 5 min at 4°C. The plate was flicked to remove the supernatant, and the pellet was resuspended in 100 µL of Fixable Viability Stain 440UV (BD Horizon™, Cat no: 566332) and incubated in the dark on ice for 15 min. After incubation, the cells were washed with Fluorescence-Activated Cell Sorting buffer (FACS, 2% FBS in PBS) and then incubated in 50 µL of primary antibody mix for 30 min in the dark at 4°C. The cells were fixed and stained for intracellular proteins using the eBioScience^TM^ Foxp3/Transcription Factor Fixation/ Permeabilization Kit (Invitrogen™, Cat no: 00-5521-00). The cells were then resuspended in 100 µL of perm fix concentrate and incubated for 30 minutes at 4°C. The cells were then washed with the perm buffer, resuspended in 50 µL of the intracellular primary antibody mix and incubated overnight at 4 °C. The next day, the cells were washed and resuspended in 200 µL of FACS buffer. The unstained and single-stain controls (for each fluorophore) were included with the samples. The single-stain controls were prepared using UltraComp eBeads™ Plus compensation beads (Invitrogen™, Cat no: 01-3333-42). The compensation beads were prepared by diluting one drop of the beads into 1 mL of FACS buffer. The bead solution was then plated in a U-bottom 96-well plate (according to the number of fluorophores being used in the panel) and centrifuged at 1500 rpm for 5 min at 4°C. The plate was then flicked to remove the supernatant, and the beads were stained with appropriate fluorophores for 30 min at 4°C. After incubation, the beads were washed and resuspended in 100 µL of FACS buffer, ready for analysis via BD FACSymphony™ A3 Cell Analyzer.

### 10x Genomics Chromium library construction and sequencing

Single-cell RNA sequencing was performed to assess gene expression and cellular heterogeneity in CD45.2⁺ immune cells isolated from mammary tumors harvested from wild-type (WT) and *Icos^-/-^* mice. Tissues were minced, digested in collagenase IV (2 mg/mL, 37 °C, 45 min), filtered, and stained with CD45.2-APCCy7 for immune cell enrichment and Sytox Green for live/dead cell discrimination. Approximately 50,000 viable CD45⁺ cells per sample were sorted on a BD FACS Aria III and processed immediately using the 10x Genomics Chromium platform for single-cell capture and cDNA preparation according to the manufacturer’s Single Cell 3’ Protocol V3.1. The silane magnetic beads and Solid Phase Reversible Immobilization (SPRI) beads were used to purify the GEM reaction mixture, and the barcoded cDNA was then amplified by PCR. The P7 and R2 primers were added during GEM incubation, and the P5 and R1 primers during library construction via end repair, A-tailing, adaptor ligation and PCR. The final libraries contain the P5 and P7 primers used in Illumina bridge amplification. Sequencing was completed on an Illumina NextSeq platform.

### Single-cell read mapping and quality control

The read information was converted to FASTQ files using bcl2fastq, mapped to the mm10-3.0.0 genome, and quantified using CellRanger (version 7.0.0) [50] with default parameters (expect-cells = 7000). The gene counts were imported to a SingleCellExperiment object [51] using DropletUtils [52] and manipulated using tidyomics [53]. Quality control and filtering were performed independently for each sample and cell type. The empty droplets were identified using barcodeRanks [52] below the inflection point. The mitochondrial gene names were downloaded using AnnotationDbi (mapIds) and EnsDb.Hsapiens.v86. Scuttle [54] was used to calculate cell quality metrics (perCellQCMetrics) using default parameters. The function isOutlier was used for each sample and cell type with default parameters to label non-viable cells based on the fraction of mitochondrial transcription. We estimated the cell cycle phase using CellCycleScoring [55]. Principal component analysis was used to verify that the cell cycle was not a major contributor to principal component loading.

For subsequent quality control and analyses, the data were converted to a Seurat container [55], and tidyseurat [56] was used for data manipulation. DoubletFinder (v3) [57] was used with default parameters to identify and filter out doublets.

### Single-cell normalisation, integration, annotation, and marker analysis

We identified variable genes within each sample using Seurat [55]. We scaled counts cell-wise and removed the unwanted variation of residual mitochondrial content using the SCTransform method [55]. The unwanted sample-wise technical variation was removed using the anchor-based integration implemented in Seurat [58]. A nearest-neighbour graph was calculated from integrated RNA abundance values, and cells were clustered using the Leiden algorithm [59]. For the first instance classification, cell-type annotations were transferred from the MouseRNAseqData [60] using SingleR [61].

We then isolated the macro clusters lymphoid, myeloid, and B-cells for independent integration, clustering, and marker gene selection. For each macro cluster, Seurat’s FindAllMarkers was used to identify cluster marker genes (with the parameters only.pos = TRUE, min.pct = 0.25, thresh.use = 0.25). The curated cell annotation was based on a consensus between the identified markers and the label transfer described above (using pbmc_multimodal as the reference). Marker genes for subclusters within each macro cluster were again identified using Seurat’s FindAllMarkers. The top 20 markers unique for each subcluster were visualised using tidyHeatmap, with both cells and features ordered to minimise Euclidean distance [62] [63]. Ribosomal features, if any, were excluded from this visualisation. t-SNE dimensionality reduction was used to visualise cell-type annotations and marker abundance [64].

### Human scRNAseq public data analysis

Raw count matrices were obtained from the integrated Human Breast Cell Atlas (iHBCA) [27], via CELLxGENE. Cell-type annotations (Tregs, CD4⁺ T cell subsets, CD8⁺ T cell subsets, NK cells) were used as provided in the published dataset. ICOS positivity was defined as raw count > 0 per cell. Per-donor ICOS⁺ cell percentages were computed within each annotated cell type and tissue compartment (healthy blood, healthy breast, ER⁺, HER2⁺, TNBC). Statistical comparisons across compartments used the Kruskal-Wallis test with Dunn’s post hoc correction; pairwise comparisons within CD4⁺ and CD8⁺ T cell subsets used Wilcoxon rank-sum tests with Benjamini-Hochberg FDR correction. Co-occurrence of ICOS⁺ Tregs with other immune cell types was quantified as Spearman’s ρ between per-donor cell-type fractions (n = 118 donors); significance was assessed by permutation test with FDR correction. Correlation of ICOS⁺ Treg fraction with CD8⁺ T cell exhaustion and effector scores (n = 90 donors) used Spearman’s ρ; scores were computed as mean normalised expression of marker gene sets (exhaustion: PDCD1, LAG3, HAVCR2, TIGIT, TOX, ENTPD1; effector: GZMB, PRF1, IFNG, GNLY).

Imaging mass cytometry data from the NeoTRIP trial (n = 1,359 ROIs) were obtained from the published dataset [28]. ICOS-high and ICOS-low ROIs were defined by a median split at the 93rd centile of the global ICOS distribution per ROI. Differential cell-type abundance between ICOS-high and ICOS-low ROIs was modelled using [29] (v2.1.34) Bayesian compositional analysis. Effect sizes are reported on the logit scale; 95% credible intervals are shown. FDR < 0.05 was considered significant.

Suppressive marker expression was compared between ICOS⁺ and ICOS⁻ Tregs (n = 19,802 cells) using paired Wilcoxon tests. Associations between the ICOS⁺ Treg fraction and the frequencies of PD-L1⁺, IDO1⁺, and PD-L1⁺IDO1⁺ APCs were assessed using Spearman’s ρ.

### Histological examination of murine tissues

For histological analysis, tissues were fixed in 10% neutral buffered formalin for 24 h at room temperature, transferred to 70% ethanol, paraffin-embedded, and sectioned at 4 µm thickness using a Leica RM 2245 microtome. Lung metastases were identified by hematoxylin and eosin (H&E) staining, in which sections were dewaxed in xylene, rehydrated through graded ethanol (100%, 90%, 70%), and stained with Mayer’s hematoxylin followed by bluing in Scott’s reagent and eosin counterstaining. Sections were then dehydrated, cleared in xylene, and mounted with Entellan medium (Sigma-Aldrich, Cat no: 1079600500) for microscopic assessment of metastatic incidence using Leica Aperio AT2 Digital Pathology Whole Slide Imaging Scanner

### Multiplex-immunohistochemistry

Breast tumor sections were stained with antibodies using the Opal 7-colour multiplex immunohistochemistry kit (Akoya Biosciences, Cat no: NEL811001KT). Staining was performed manually per the manufacturer’s protocol, as previously described (79). The following antibodies were used: CD3e (Clone: SP7, Dilution 1:150, Invitrogen™, Cat no: MA1-90582), Anti-pan Cytokeratin (Clone: AE1/AE3+5D3, Dilution 1:200, Abcam, Cat no: ab86734), FoxP3 (Clone: 236A/E7, Dilution 1:300, Abcam, Cat no: ab20034) CD8 (Clone: 4B11, Dilution 1:500, Invitrogen™, Cat no: MA1-80231), ICOS (Clone: SP98, Dilution 1:400 Invitrogen™, Cat no: ab105227), and spectral DAPI (Akoya Biosciences). Whole-slide scans at 10x magnification were performed using the Vectra 3 Automated Quantitative Pathology Imaging System (PerkinElmer) to mark regions of interest, followed by acquisition of 20x magnification multispectral images (MSI) for analysis. A spectral library was created on inForm Automated Image Analysis Software (Akoya Biosciences) to analyse cell populations quantitatively. Immune cell populations of interest were identified by each marker’s mean fluorescence intensity (MFI), and the algorithm was applied to all samples. Images were analysed on the HALO® image analysis platform (Indica Labs) using the Highplex v4.1.3 module for target cell phenotyping.

### *In vitro* PMA/Ionomycin assay

PMA and Ionomycin stimulation were conducted to activate transcription factors involved in intracellular signalling and to induce cytokine production in various types of immune cells ex vivo. Before the antibody staining for flow cytometry, the immune cells isolated from different tissue types were transferred to a U-bottom 96-well plate and centrifuged at 1700 rpm for 5 min at room temperature. The supernatant was removed, and the cells were resuspended in 100 µL of restimulation cocktail containing 5 ng/mL PMA, 500 ng/mL ionomycin, 1 µL/mL Brefeldin A (BD GolgiPlug™, Cat no: 555029) in Complete Iscove modified Dulbecco media (cIMDM). The cIMDM was prepared by supplementing IMDM (Gibco™, Cat no: 12440053) with 10% (vol/vol) heat-inactivated FBS, 100 U/mL penicillin-streptomycin, 2 mmol/L L-glutamine, 1 mmol/L sodium pyruvate, and 55 μmol/L 2-mercaptoethanol. Cells were incubated for 3-4 hours at 37°C, 5% CO_2_. After incubation, the cells were washed with pre-warmed cIMDM media and prepared for flow cytometry staining, as mentioned above.

### Whole-mount staining of mammary glands

To visualise the structure of the mammary gland at different developmental stages, the mammary glands were carefully excised and mounted on microscope slides. A carmine alum solution containing 1 g of carmine and 2.5 g of potassium sulphate was prepared by diluting 500 mL of distilled water with 1 g of Carmine (Sigma-Aldrich, Cat no: C1022) and 2.5 g of potassium sulphate 1 day before dissection. The solution was then boiled for 20 min in a 1 L conical flask, and the final volume was adjusted to 500 mL with MilliQ water. The tissue slides are fixed in Carnoy’s fixative (100% ethanol, chloroform, and glacial acetic acid in a 6:3:1 ratio) for 4 hours at RT or overnight at 4°C. The slides were washed with 70% ethanol for 15 min, then rehydrated in 35% and 15% ethanol for 5 min each, followed by a 5 min wash in distilled water. The slides were incubated in carmine alum solution overnight at RT. To clear the tissue, the carmine alum-stained slides were incubated in serial ethanol baths (50%, 70%, 95%, 100%) for 5 min each and incubated in xylene overnight at room temperature. Once the tissue was cleared, an Entellan mounting medium was added to the slides to mount the coverslip. Whole slide scans were obtained using the Leica Aperio AT2 Digital Pathology Whole Slide Imaging Scanner.

### RNA isolation, cDNA synthesis and qPCR

Total RNA was isolated from cancer cell pellets or snap-frozen tumor tissues using the Qiagen RNeasy Mini Kit (QIAGEN, Cat no: 74104) following the manufacturer’s instructions. Briefly, cell pellets or homogenised tissue were lysed in Qiazol reagent, followed by chloroform extraction and ethanol precipitation before loading onto RNeasy columns. On-column DNase (QIAGEN, Cat no: 79254) digestion was performed to remove genomic DNA, and RNA was eluted in RNase-free water (QIAGEN, Cat no: 129112). RNA integrity and concentration were assessed by 4200 TapeStation System (Agilent). For cDNA synthesis, RNA was reverse-transcribed using oligo(dT) primers and dNTPs in a thermocycler, and cDNA was stored at -20°C. Real-time quantitative PCR (qPCR) was performed using β-actin as the housekeeping gene. Reactions were run in triplicate on a ViiA 7 Real-Time PCR System (Applied Biosystems), with the cycling program consisting of enzyme activation (95°C, 20 sec) followed by 40 cycles of denaturation (95°C, 1 sec) and annealing/extension (60°C, 20-60 sec). Relative gene expression was calculated using the 2^-ΔΔCT^ method.

### Data availability

The single-cell RNA sequencing data generated in this study are available in the Gene Expression Omnibus (GEO) under accession number GSE348069. The publicly available datasets analyzed in this study were obtained from the integrated Human Breast Cell Atlas (iHBCA) via CELLxGENE at https://cellxgene.cziscience.com/collections/48259aa8-f168-4bf5-b797-af8e88da6637 (dataset “Integrated HBCA”) and from the NeoTRIPaPDL1 imaging mass cytometry dataset [28], available through Zenodo at https://doi.org/10.5281/zenodo.7990870. All other data generated in this study, including flow cytometry, multiplex immunohistochemistry, qPCR, and histology data, are available from the corresponding author upon request.

