## Supplemental Files for "ICOS blockade alleviates immune suppression in breast tumor microenvironment"

#### **Supplemental Information**

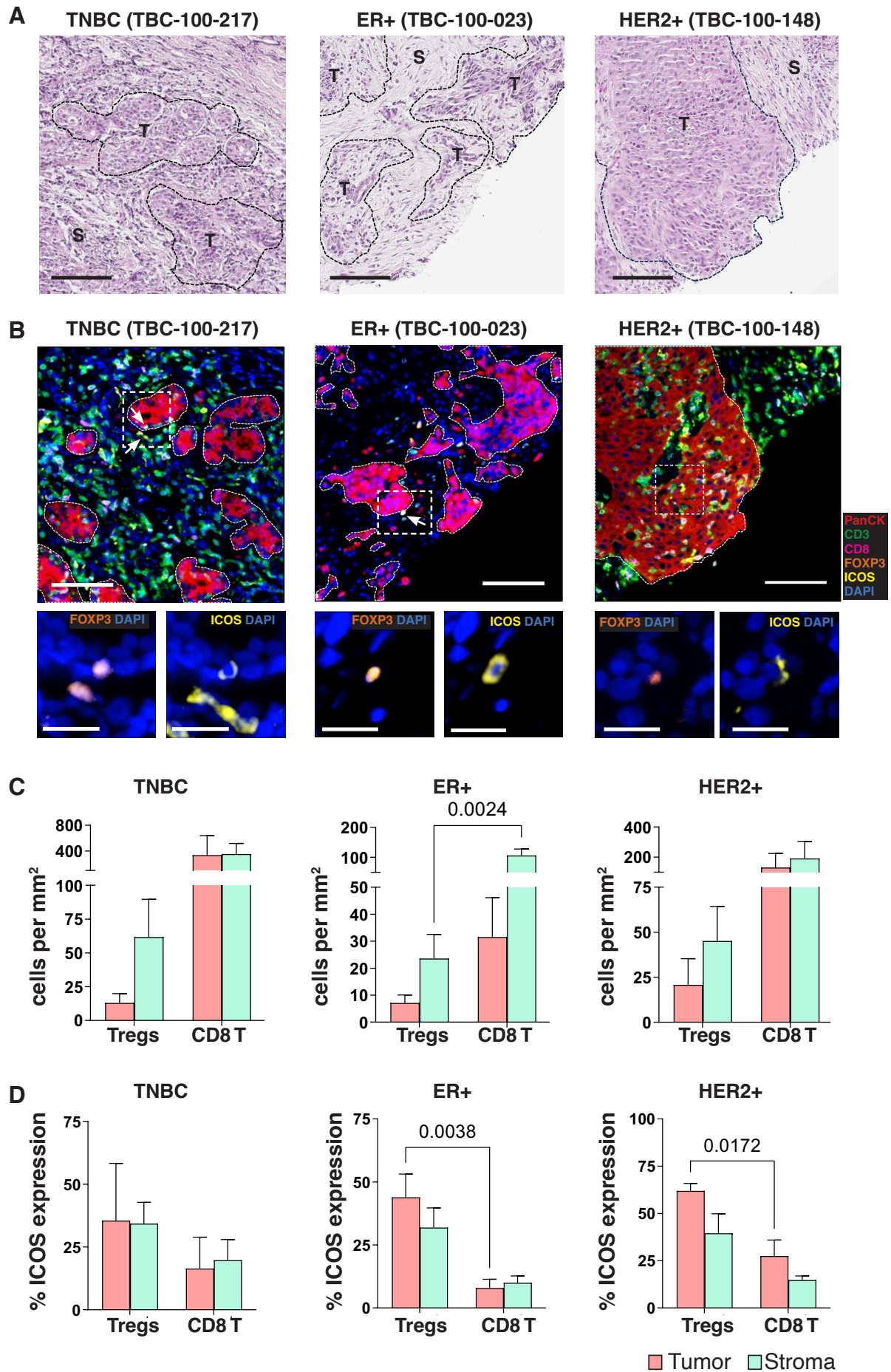

**Supplementary Figure 1. ICOS<sup>+</sup> Tregs are present across breast cancer subtypes. A.**

Representative hematoxylin and eosin-stained (H&E) sections of human breast cancer specimens from triple-negative breast cancer (TNBC), estrogen receptor-positive (ER<sup>+</sup>), and HER2-positive (HER2<sup>+</sup>) subtypes. Tumor (T) and stromal (S) regions are delineated by dashed lines. **B.** Representative multiplex immunofluorescence images of breast cancer specimens from TNBC, ER<sup>+</sup>, and HER2<sup>+</sup> subtypes showing spatial localization of immune cells within the tumor compartment. Pan-cytokeratin (PanCK, red) marks tumor epithelial cells. CD3 (green) identifies total T cells. CD8 (magenta) identifies cytotoxic T cells. FOXP3 (orange) marks regulatory T cells. ICOS (yellow) marks ICOS-expressing immune cells. DAPI (blue) labels nuclei. Dashed box indicates region of interest shown at higher magnification in adjacent panels, highlighting FOXP3<sup>+</sup> and ICOS<sup>+</sup> cells (arrows). Scale bar, 100 μm. **C.** Quantification of Tregs and CD8<sup>+</sup> T cells (cells per mm<sup>2</sup>) within tumor (pink) and stromal (green) compartments across TNBC, ER<sup>+</sup>, and HER2<sup>+</sup> breast cancers. **D.** Percentage of ICOS-expressing cells within Treg and CD8<sup>+</sup> T-cell populations in tumor and stromal regions for each subtype. Data are presented as mean ± s.e.m. $p < 0.05$ was considered significant.

# A

#### ICOS+ vs ICOS- Treg suppressive marker expression (n=19,802 Tregs)

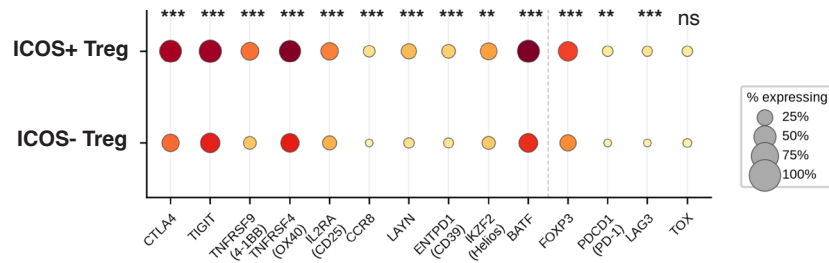

# B

#### ICOS+ Treg burden correlates with immunosuppressive APC frequency

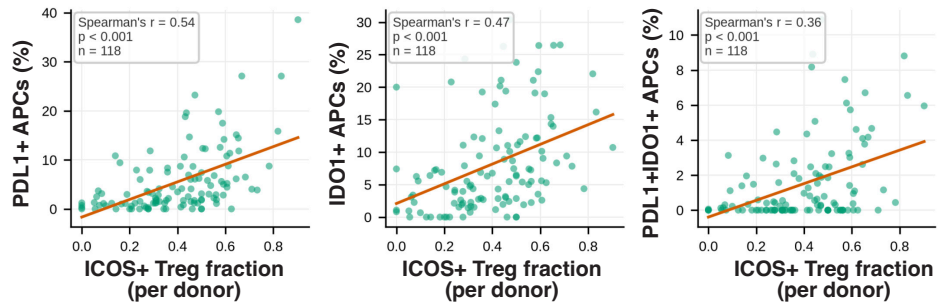

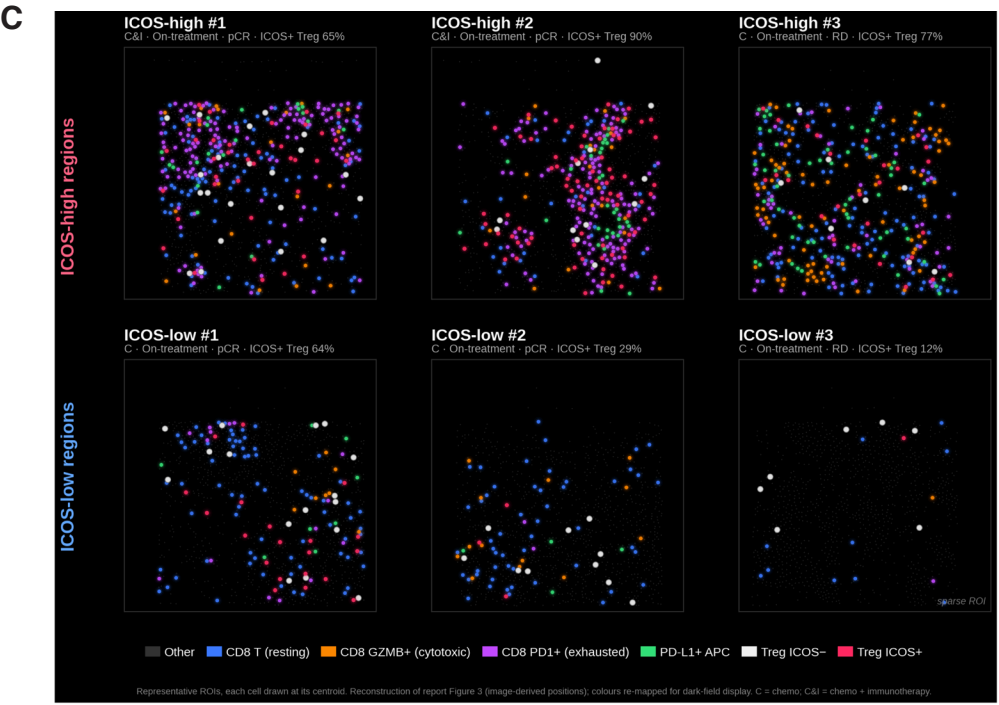

# D

| Arm | Outcome | n ROIs | ICOS<sup>-</sup> Treg median | ICOS<sup>+</sup> Treg median | $\Delta$ (ICOS<sup>-</sup> - ICOS<sup>+</sup>) | p-value (paired Wilcoxon) | Sig. |
| --- | --- | --- | --- | --- | --- | --- | --- |
| Chemo (C) | RD | 115 | 0.099 | 0.124 | -0.016 | 0.0103 | * |
| Chemo (C) | pCR | 92 | 0.120 | 0.134 | -0.015 | 0.0117 | * |
| Chemo + Atezo (C&I) | RD | 114 | 0.098 | 0.100 | -0.001 | 0.7880 | ns |
| Chemo + Atezo (C&I) | pCR | 95 | 0.133 | 0.131 | +0.002 | 0.7480 | ns |

# E

| Arm | Outcome | n ROIs | ICOS<sup>-</sup> Treg exhausted share | ICOS<sup>+</sup> Treg exhausted share | p-value (paired Wilcoxon) | Sig. |
| --- | --- | --- | --- | --- | --- | --- |
| Chemo (C) | RD | 101 | 23.2% | 18.6% | 0.01010 | * |
| Chemo (C) | pCR | 88 | 22.7% | 20.0% | 0.16400 | ns |
| Chemo + Atezo (C&I) | RD | 93 | 28.6% | 19.6% | 0.00399 | ** |
| Chemo + Atezo (C&I) | pCR | 86 | 24.0% | 16.7% | 0.07040 | ns |

**Supplementary Figure 2. ICOS<sup>+</sup> Treg phenotype, spatial co-localization, and CD8<sup>+</sup> T cell neighbor analysis.** **A.** Dot plot comparing suppressive marker expression between ICOS<sup>+</sup> and ICOS<sup>-</sup> Tregs (n=19,802 Tregs, iHBCA). Dot size encodes the percentage of cells expressing each marker. Color encodes mean expression (log-normalized counts). Markers left of the dashed divider comprise the 10-gene tumor-resident Treg signature: CTLA4, TIGIT, TNFRSF9 (4-1BB), TNFRSF4 (OX40), IL2RA (CD25), CCR8, LAYN, ENTPD1 (CD39), IKZF2 (Helios), BATF; markers right of the divider are FOXP3, PDCD1 (PD-1), LAG3 and TOX. Per-donor paired Wilcoxon test; \*\*\*, p<0.001; \*\*, p<0.01; ns, not significant. **B.** Scatter plots showing Spearman correlations between ICOS<sup>+</sup> Treg fraction and the frequency of PD-L1<sup>+</sup> APCs (left; $\rho=0.54$, p<0.001), IDO1<sup>+</sup> APCs (center; $\rho=0.47$, p<0.001), and PD-L1<sup>+</sup>IDO1<sup>+</sup> APCs (right; $\rho=0.36$, p<0.001) across n=118 donors (iHBCA). Orange line, linear regression fit. **C.** Fluorescence-style spatial maps of immune cell centroids from representative NeoTRIP IMC ROIs, three ICOS-high ROIs (top row) and three ICOS-low ROIs (bottom row). Each dot represents a single cell centroid. Color key: Other (grey), CD8<sup>+</sup> T resting (blue), CD8<sup>+</sup>GZMB<sup>+</sup> T (orange), CD8<sup>+</sup>PD1<sup>+</sup> T exhausted (purple), PD-L1<sup>+</sup> APC (green), ICOS<sup>-</sup> Treg (white), ICOS<sup>+</sup> Treg (pink). ICOS-high/low was defined as in Figure 1D. **D.** CD8<sup>+</sup> T cell neighbor fraction for ICOS<sup>+</sup> vs ICOS<sup>-</sup> Tregs across NeoTRIP IMC ROIs, stratified by treatment arm (Chemo [C] and Chemo + Atezolizumab [C&I]) and pathological response (RD, residual disease, and pCR, pathological complete response). Median CD8<sup>+</sup> neighbor fraction per Treg is shown. $\Delta$ = ICOS<sup>+</sup> - ICOS<sup>-</sup> median. Paired Wilcoxon test within each arm × outcome stratum; \*, p<0.05; ns, not significant. **E.** Exhausted CD8<sup>+</sup> T cell share among Treg neighbors (CD8<sup>+</sup>PD1<sup>+</sup>T\_Ex / all CD8<sup>+</sup> Treg neighbors), comparing ICOS<sup>+</sup> vs ICOS<sup>-</sup> Tregs across the same strata as in D (NeoTRIP IMC). Paired Wilcoxon test; \*, p<0.05; \*\*, p<0.01; ns, not significant.

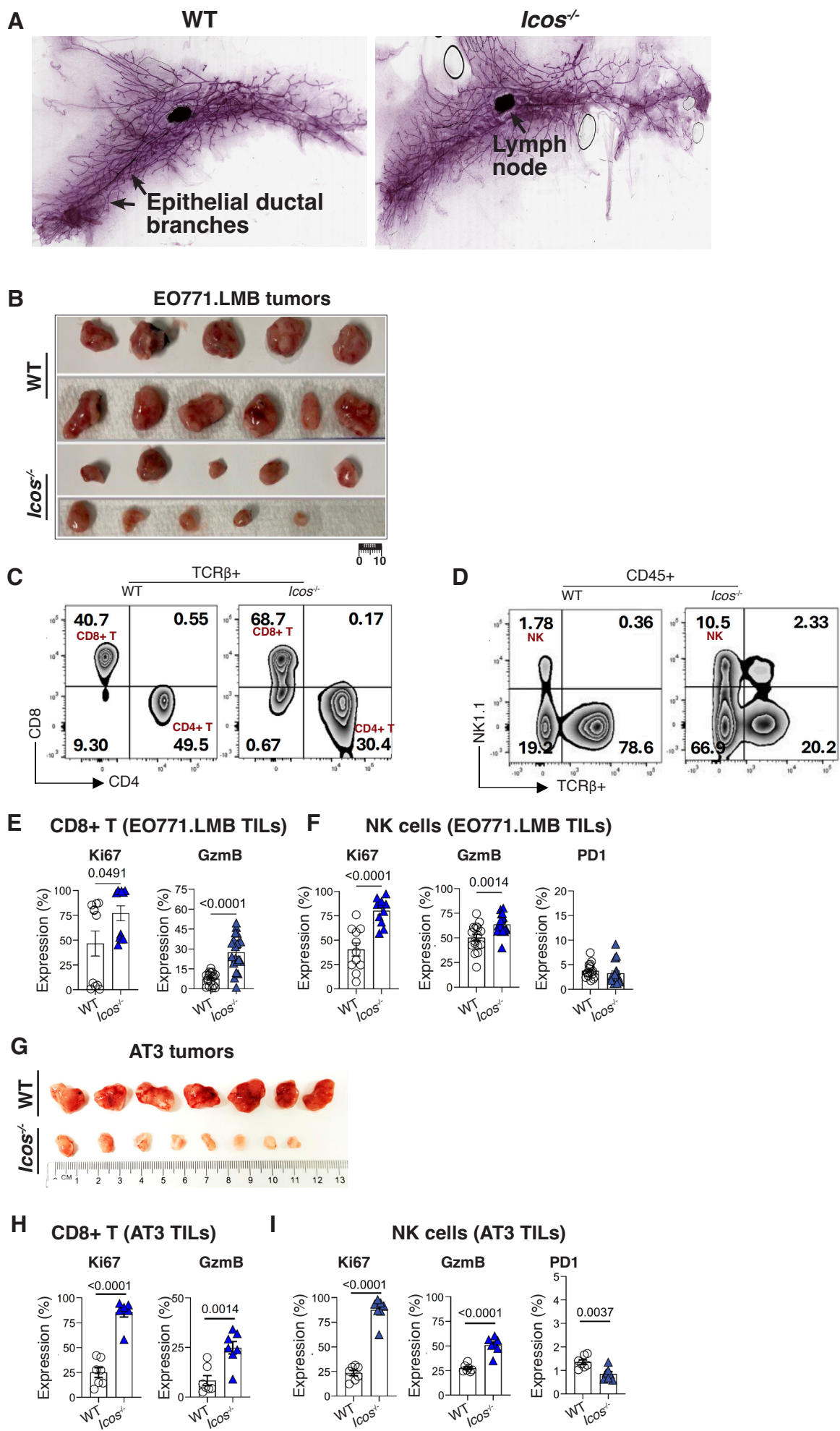

**Supplementary Figure 3. ICOS deficiency does not alter pubertal mammary gland development but impairs mammary tumor growth.** **A.** Representative whole-mount carmine alum staining of mammary glands from wild-type (WT) and *Icos*<sup>-/-</sup> C57BL/6 female mice (8-12 weeks), showing comparable epithelial ductal branching and overall mammary gland architecture. Epithelial ductal branches and mammary lymph nodes are indicated by arrows. **B.** Images of EO771.LMB mammary tumors harvested from WT and *Icos*<sup>-/-</sup> mice at the experimental endpoint. Scale bar, 10 mm. **C.** Representative flow cytometry plots of TCRβ<sup>+</sup> T cells from EO771.LMB tumors, showing CD8<sup>+</sup> T-cell and CD4<sup>+</sup> T-cell compartments in WT control and *Icos*<sup>-/-</sup> mice. **D.** Representative flow cytometry plots of CD45<sup>+</sup> tumor-infiltrating immune cells from EO771.LMB tumors, showing NK cell frequency in *Icos*<sup>-/-</sup> tumors compared with WT controls. **E.** Flow cytometric quantification of Ki67 and GzmB expression in tumor-infiltrating CD8<sup>+</sup> T cells from EO771.LMB tumors. **F.** Flow cytometric quantification of Ki67, GzmB, and PD-1 expression in tumor-infiltrating NK cells from EO771.LMB tumors. **G.** Images of AT3 mammary tumors harvested from WT and *Icos*<sup>-/-</sup> mice at the experimental endpoint. Scale bar, 10 mm. **H.** Flow cytometric quantification of Ki67 and GzmB expression in tumor-infiltrating CD8<sup>+</sup> T cells from AT3 tumors. **I.** Flow cytometric quantification of Ki67, GzmB, and PD-1 expression in tumor-infiltrating NK cells from AT3 tumors. Data are presented as mean ± s.e.m. Statistical significance was determined using unpaired two-tailed t-tests. p<0.05 was considered significant.

Top marker genes Tregs ( WT vs *Icos*<sup>-/-</sup>)

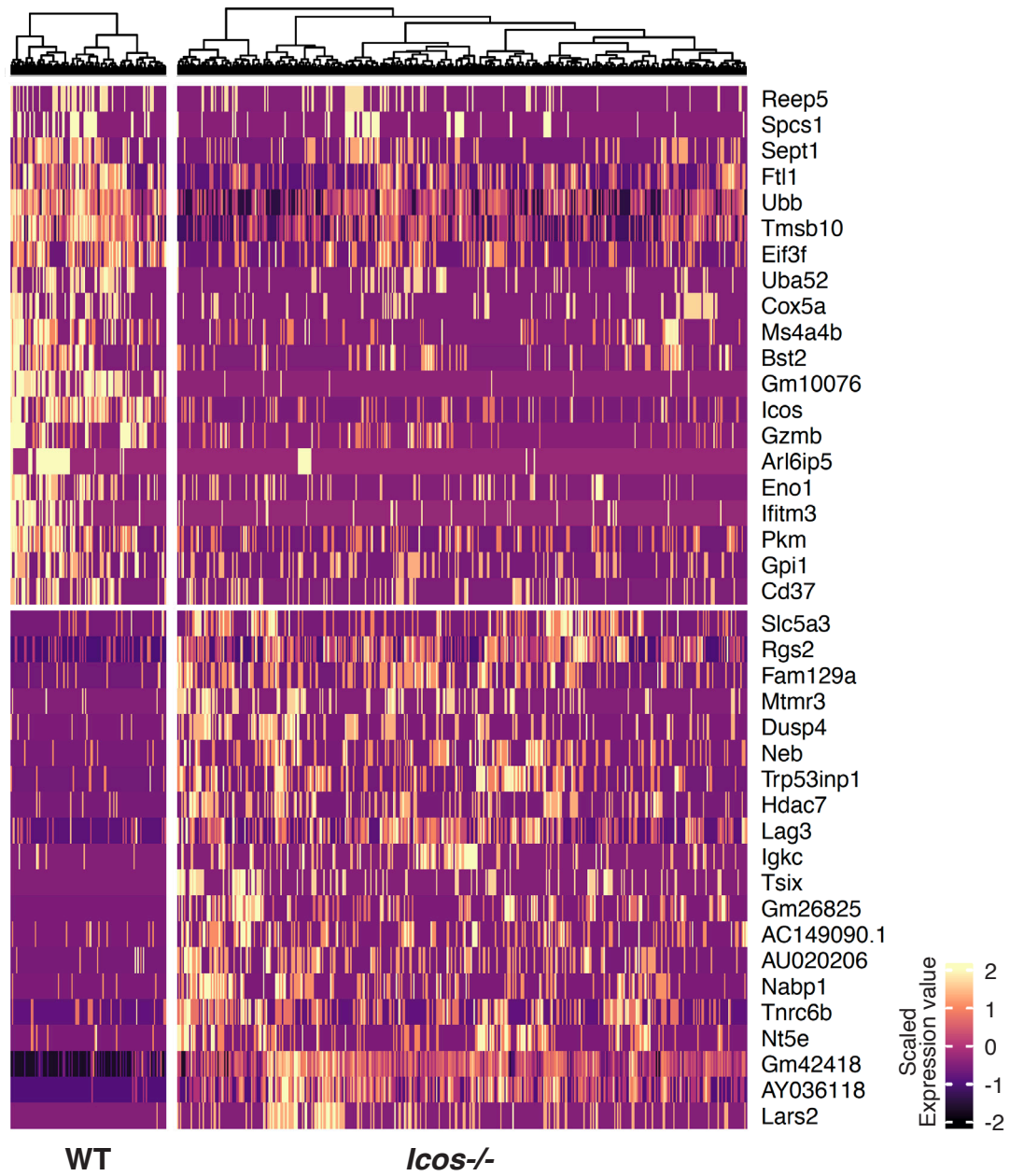

**Supplementary Figure 4. Gene expression analysis of Tregs within WT vs *Icos*<sup>-/-</sup> TILs.**

Heatmap showing the transcript abundance of the top marker genes distinguishing WT from *Icos*<sup>-/-</sup> Tregs by single-cell RNA sequencing. Gene abundances are displayed as scaled SCT values.

### Top marker genes - CD8+ T cell subsets

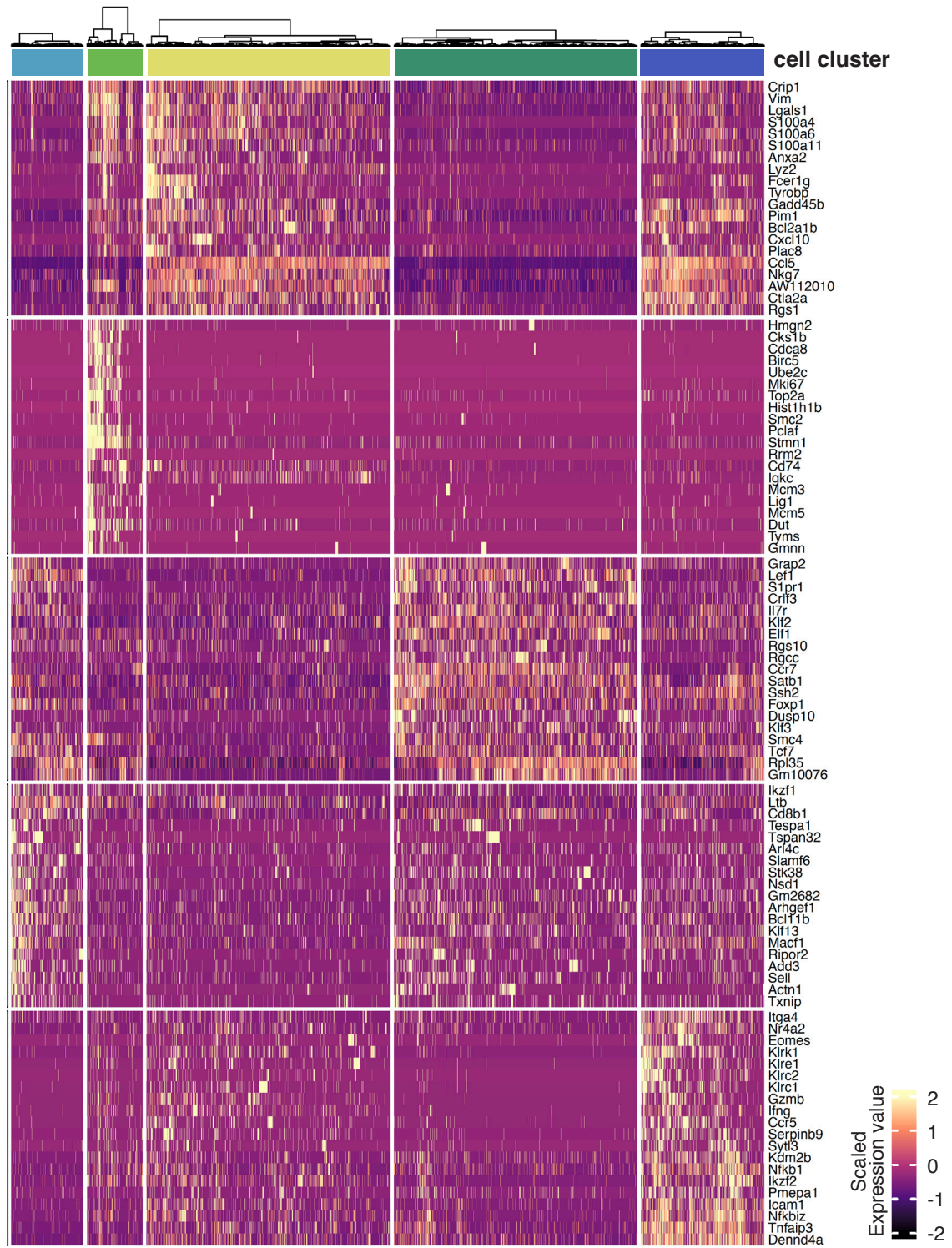

**Supplementary Figure 5. Gene expression analysis of CD8<sup>+</sup> T cell subsets.** Heatmap depicting the scaled transcript abundance of the top subset-defining marker genes across reclustered CD8<sup>+</sup> T cell populations identified by single-cell RNA sequencing. Cell cluster color key corresponds to the CD8<sup>+</sup> T cell clusters shown in Fig. 3C. Gene expression levels are displayed as scaled SCT-normalized values.

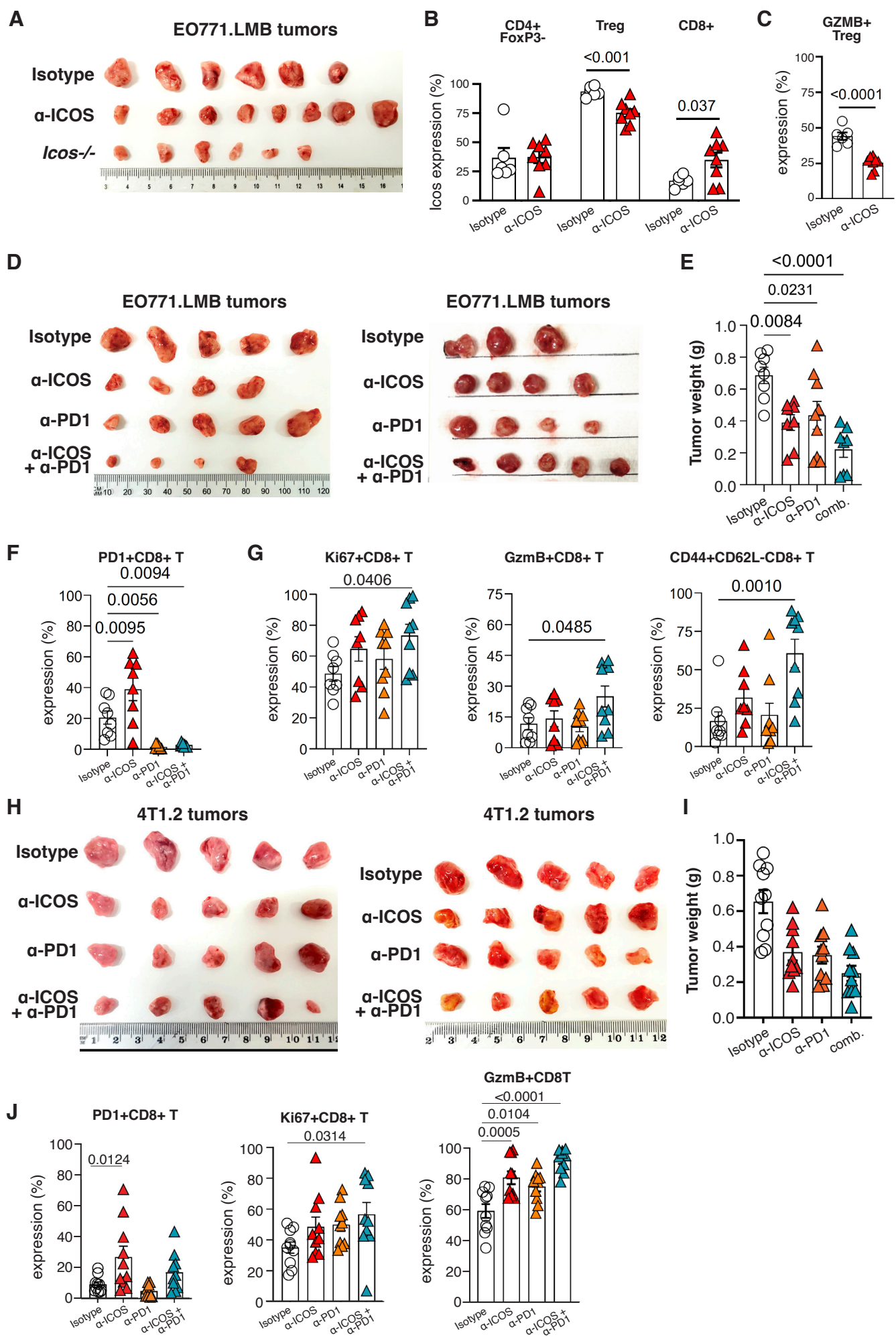

**Supplementary Figure 6. ICOS blockade reduces tumor burden across multiple breast cancer models.** **A.** EO771.LMB mammary tumors isolated from mice treated with isotype control or anti-ICOS antibodies. The last row corresponds to tumors grown in *Icos*<sup>-/-</sup> female mice as a control. **B.** Flow cytometric quantification of ICOS expression on intratumoral FOXP3<sup>-</sup>CD4<sup>+</sup> T cells, Tregs, and CD8<sup>+</sup> T cells in EO771.LMB tumors from mice treated with isotype control or anti-ICOS antibodies, confirming on-target antibody activity. **C.** Frequency of GzmB<sup>+</sup> Tregs in EO771.LMB tumors following anti-ICOS treatment. **D.** Images of EO771.LMB tumors harvested at the experimental endpoint from mice treated with isotype control, anti-ICOS, anti-PD-1, or the combination of anti-ICOS and anti-PD-1 antibodies. **E.** Endpoint tumor weight of EO771.LMB tumors across treatment groups. **F.** Quantification of PD-1<sup>+</sup> CD8<sup>+</sup> T cells in EO771.LMB tumors across treatment groups. **G.** Quantification of Ki67<sup>+</sup>, GzmB<sup>+</sup>, and CD44<sup>+</sup>CD62L<sup>-</sup> effector-memory CD8<sup>+</sup> T cells in EO771.LMB tumors across treatment groups. **H.** Images of 4T1.2 mammary tumors harvested at the experimental endpoint across treatment groups. **I.** 4T1.2 tumor weight across treatment groups. **J.** Quantification of PD-1<sup>+</sup>, Ki67<sup>+</sup>, and GzmB<sup>+</sup> CD8<sup>+</sup> T cells in 4T1.2 tumors across treatment groups. Data are presented as mean ± s.e.m. Statistical significance was determined using unpaired two-tailed t-tests, with P values indicated in the plots.

**Supplementary Table 1:** List of breast tumor samples used for mIHC

|  | **Patient ID** | **Age at**<br>**Diagnosis** |
| --- | --- | --- |
| **ER+** | TBC-100-222 | 45 |
|  | TBC-100-163 | 44 |
|  | TBC-100-076 | 28 |
|  | TBC-100-023 | 64 |
| **HER2+** | TBC-100-228 | 35 |
|  | TBC-100-148 | 52 |
|  | TBC-100-140 | 79 |
|  | TBC-100-079 | 58 |
| **TNBC** | TBC-100-217 | 49 |
|  | TBC-100-109 | 42 |
|  | TBC-100-052 | 34 |

**Supplementary Table 2:** List of antibodies used for flow cytometry

| **Antibody** | **Fluorophore** | **Clone** | **Isotype** | **Company** | **Catalogue No.** |
| --- | --- | --- | --- | --- | --- |
| CD45.2 | Alexa 700 | 104 | Mouse IgG2a, k | Thermofisher | 56-0454-8 2 |
| CD45.2 | APC/Cyanine7 | 30-F11 | Rat IgG2b, k | BioLegend | 103116 |
| NK1.1 | BUV805 | PK136 | Mouse IgG2a, k | BD Bioscience | 741926 |
| NKp46 | R718 | 29A1.4 (RUO) | Rat IgG2b, k | BD Bioscience | 751865 |
| TCRβ | BUV496 | H57-597 | Armenian Hamster IgG2, λ1 | BD Bioscience | 749915 |
| CD4 | BUV395 | GK1.5 | Rat IgG2b, k | BD Bioscience | 563790 |
| CD8a | BV570 | 53-6.7 | Rat IgG2a, k | BioLegend | 100739 |
| Epcam | PE | G8.8 | Rat IgG2a, k | Thermofisher | 12-5791-82 |
| Foxp3 | FITC | FJK-16s | Rat IgG2a k | Thermofisher | 11-5773-82 |
| Foxp3 | BV-421 | MF-14 | Rat IgG2a k | BioLegend | 126419 |
| Foxp3 | PE | FJK-16s | Rat IgG2a k | Thermofisher | 12-5773-82 |
| CD25 | BV510 | PC61 | Rat IgG2, λ1 | BD Bioscience | 566202 |
| PD-1 | mCherry | J43 | Armenian Hamster IgG2, $\kappa$ | BD Bioscience | 562523 |
| Ki67 | BV605 | 16A8 | Rat IgG2a, k | BioLegend | 652413 |
| KLRG1 | BV711/<br>BUV615 | 2F1 | Syrian Hamster IgG2, $\kappa$ | BD Bioscience | 564014 |
| CD62L | BUV737 | MEL-14 | Rat IgG2a, k | BD Bioscience | 612833 |
| LAG-3 | PerCP.Cy5.5 | C9B7W | Council IgG1, k | BioLegend | 125211 |

| Antibody | Fluorophore | Clone | Isotype | Company | Catalogue No. |
| --- | --- | --- | --- | --- | --- |
| ICOS | BV421 | C398.4A | American Hamster IgG | BioLegend | 313523 |
| CD44 | PE | IM7.8.1 | Rat IgG2b | Thermofisher | MA5-17880 |
| TIM-3 | PE/Cyanine7 | RMT3-23 | Rat IgG2a, $\kappa$ | BioLegend | 119716 |
| Granzyme B(GzmB) | APC | GB11 | Mouse IgG1, $\kappa$ | BioLegend | 515405 |
| IFNγ | BV785 | XMG1.2 | Rat IgG1, $\kappa$ | BD Bioscience | 566366 |
| TNF-α | BV650 | MP6-XT22 | Rat IgG2a, $\kappa$ | BioLegend | 506333 |
| CD69 | PE/Cyanine5 | H1.2F3 | American Hamster IgG | BioLegend | 104509 |
| CD103 | BV711 | 2 E7 | American Hamster IgG | BioLegend | 121435 |
| Fixable Viability Dye | UV 440 | NA | NA | BD Bioscience | 566332 |
